# Binding the acoustic features of an auditory source through temporal coherence

**DOI:** 10.1101/2021.05.05.442748

**Authors:** Mohsen Rezaeizadeh, Shihab Shamma

## Abstract

Numerous studies have suggested that the perception of a target sound source can only be segregated from a complex acoustic background if the acoustic features underlying its perceptual attributes (e.g., pitch, location, and timbre) induce temporally modulated responses that are mutually correlated, and that are uncorrelated from those of other sources in the mixture. This “temporal coherence” hypothesis asserts that listening attentively to one or a subset of attributes of a target source enhances their neural responses and concomitantly enhances all other coherent responses, thus binding them together while simultaneously suppressing the incoherent responses to the background features. Here we report on EEG measurements in human subjects engaged in various sound segregation tasks that demonstrate rapid binding among the temporally coherent features of the attended source regardless of their identity, harmonic relationship, or frequency separation, thus confirming the key role temporal coherence plays in the organization of auditory scenes.

## Introduction

Humans and other animals can segregate a target sound from background interference and noise with remarkable ease (***Bregman, 1990***; ***Moore and Gockel, 2002***; ***Middlebrooks et al., 2017***), despite the highly interleaved spectrotemporal acoustic components of the different sound sources (or streams) (***Brungart et al., 2001***). It is hypothesized that attention is important for this process to occur in a listener’s brain, and that the consistent or *coherent* temporal co-modulation of the acoustic features of the target sound, and their incoherence from those of other sources, are the two key factors that induce the binding of the target features and its emergence as the foreground sound source (***Lu et al., 2017***; ***Shamma et al., 2011***; ***Elhilali et al., 2009***). Specifically, the *temporal coherence principle* implies that acoustic features underlying the perceptual attributes of a sound emanating from a single source (e.g., its pitch, timbre, location, loudness) evoke correlated neural responses, i.e., that fluctuate similarly in power over time, and that the attentive listener tracks and utilizes this neural coherence to extract and perceive the source. The definition of temporal coherence and other related terms is further elaborated upon in the Methods ***section***.

Numerous studies have provided insights into the temporal coherence theory and tested its predictions. For example, psychoacoustic experiments have shown that perception of synchronous tone sequences as belonging to a single stream is not appreciably affected by their frequency separation (from 3 semitones to over an octave) or small frequency fluctuations of the individual components, as long as the tones remain temporally coherent (***Micheyl et al., 2013a***, b). Further-more, it is far easier to detect the temporal onset misalignment between tones across two synchronized sequences, compared to between asynchronous (e.g., alternating) sequences (***Elhilali et al., 2009***), suggesting that temporally coherent tone sequences are perceived as a single stream (***Bregman and Campbell, 1971***; ***Zera, 1993***; ***Zera and Green, 1995***). Additional strong evidence for the temporal coherence principle was provided by a series of experiments utilizing the stochastic figure–ground stimulus, in which synchronous tones (referred to as the “figure“) are found to pop out perceptually against a background of random desynchronized tones, with the perceptual saliency of the “figure” being proportional to the number of its coherent tones (***Teki et al., 2013***, ***2016***; ***O’Sullivan et al., 2015***).

To account for the neural bases underlying the principle of temporal coherence, a recent electrocorticography (ECoG) study in human patients examined the progressive extraction of attended speech in a multi-talker scenario. It demonstrated that a linear mapping could transform the multi-talker responses in the human primary auditory cortex (Heschl’s Gyrus, or HG, in humans) to those of the attended speaker in higher auditory areas. Furthermore, the mapping weights could be readily predicted by the mutual correlation, or *temporal correlation* between the responses in the HG sites (***O’Sullivan et al., 2019***). This experimental finding is consistent with an earlier computational model for how temporal coherence could be successfully implemented by measuring the coincidence of acoustic feature responses to perform speech segregation (***Krishnan et al., 2014***). It has also validated single-unit studies in ferret auditory cortex, which tested the importance of attention and temporal coherence in stream formation and selection, and further demonstrated that the responses and connectivity among responsive neurons were rapidly enhanced by synchronous stimuli and suppressed by asynchronous sounds, but only when the ferrets actively attended to the stimuli (***Lu et al., 2017***). Exactly the same idea has been shown to be relevant in the binding of multisensory auditory-visual streams both in cortical responses and in psychoacoustic tests (***Bizley et al., 2016***; ***Atilgan et al., 2018***; ***Atilgan and Bizley, 2021***), as well as to explain stream formation associated with comodulation masking release (***Krogholt Christiansen and Oxenham, 2014***)

In this study, we sought to investigate the properties and dynamics of the temporal coherence principle using the more accessible EEG recordings in human subjects while performing psychoacoustic tasks with a wide variety of stimuli, including natural speech. The experiments tested several key predictions of the temporal coherence hypothesis (schematized in ***Figure 1***), primarily the coincidence of the neural responses to any acoustic features is the fundamental and overriding determinant of the segregated perception of an auditory stream. Thus, it is *not* the specific nature of the features (e.g., being a single-tone, tone-complex, or a noise burst) or the harmonic relationship among the tones of the complex that determines their binding. Rather, it is the temporal coincidence among the components that matter. A second prediction of the hypothesis is that directing attention to a specific feature (e.g., a tone in a complex) not only enhances (or modulates) the response of the neurons tuned to it but would also bind it or similarly modulate the responses of all other neurons that are synchronized with it. Conversely, attending to a target sound is postulated to suppresses responses due to acoustic features that are highly uncorrelated with the target. Another aspect of the temporal coherence hypothesis that has already been explored is the rapid dynamic nature of the binding among the components of a stream (***Lu et al., 2017***), which explains how listeners are able to switch attention and rapidly reorganize their auditory scene according to their desired focus. Nevertheless, the role of attention in stream formation can be somewhat ambiguous in that many studies have demonstrated streaming indicators even in the absence of selective attention (***O’Sullivan et al., 2015***; ***Sussman, 2017***). However, even in these cases, deploying selective attention always enhanced the responses, significantly confirming its important role in mediating the streaming percept.

**Figure 1.**
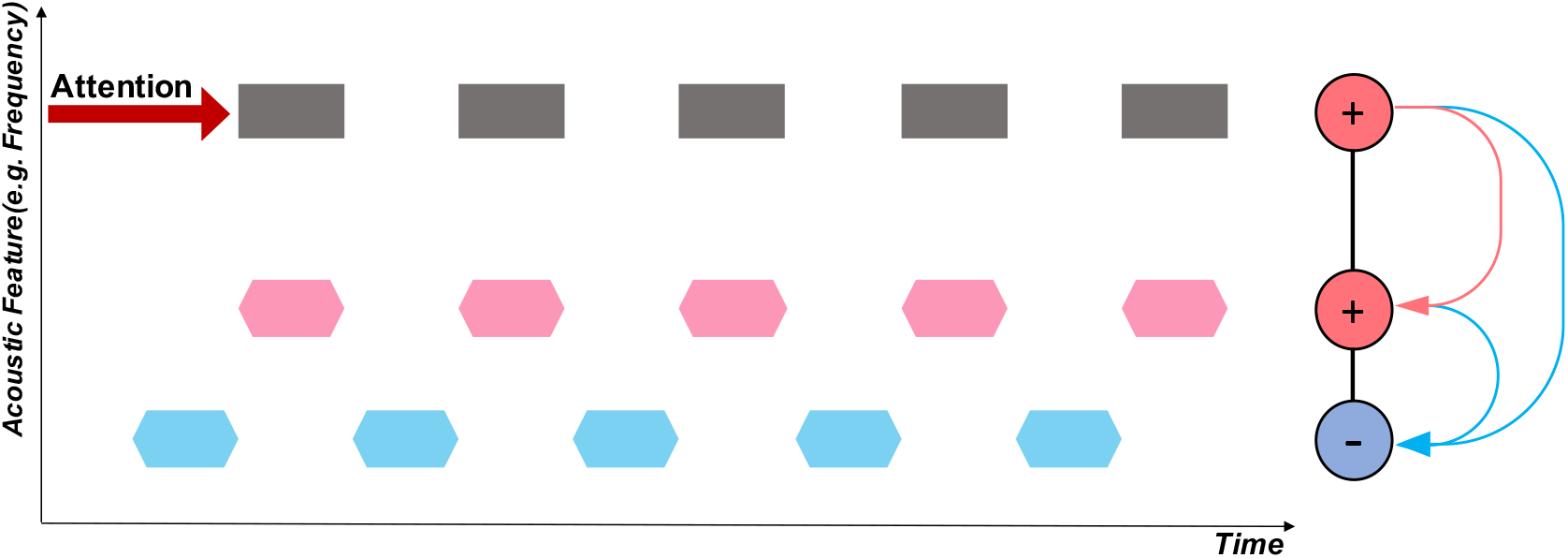
Schematic of attention and the temporal coherence principle. The horizontal axis is time, and the vertical axis depicts an arbitrary feature dimension of interest (e.g., spectral frequency, fundamental frequency (pitch), or location). Three separate sequences of sound tokens are depicted: two are synchronized (in black/pink), alternating (or de-synchronized) with another sequence (in blue). If attention is focused on the black sequence, its neural responses become enhanced. Because of its temporal coherence with the pink sequence, mutual excitation causes the two to bind, thus becoming enhanced together. By contrast, the blue stream is asynchronous (temporally incoherent), and hence it is suppressed by rapidly formed mutually inhibitory interactions (depicted in blue).

However, it should be noted that conducting these experiments and analyses with EEG recordings is difficult because of the extensive spatial spread of the responses across the scalp. This introduces two types of challenges that must be overcome. First, it is hard to resolve and assess the binding of individual frequency components in a complex or in a speech mixture that contains many other nearby components. Second, because the responses from many neural sources interact and superimpose in EEG recordings, a response enhancement due, for example, to attention to a specific feature may be accompanied by suppression of responses from a competing feature. Hence, the total response becomes instead manifested as a complex, unintuitive modulation of the response patterns. Both of these challenges can be overcome by the techniques presented in this report. Specifically, we addressed the spectral resolution problem by presenting isolated tone probes *immediately after* the end of the complex stimuli. By aligning a probe tone with various spectral components of the preceding stimulus and measuring the *persistent* effects of attention on its responses just after the stimulus, we could detect the attentional effects on the responses to individual components within these complex stimuli. Furthermore, to decode and assess these changes directly from the complicated distributed EEG responses, we resorted to a pattern classification technique that quantified whether attention significantly altered (or modulated) the response patterns and for how long before and after.

## Results

The results described here are of EEG experiments conducted on normal-hearing subjects. Details of the experimental setup, subjects, and stimuli are provided in each subsection below, as well as in the Methods ***section***. The experiments begin by exploring the consequences of temporal coherence on simple harmonic tone-complexes at a uniform rate and progress to inharmonic complexes, irregular presentation rates, mixed tone and noise sequences, and ending with speech mixtures.

Note that all of the stimulus paradigms in this study were selected to closely resemble “classical” paradigms of streaming, e.g., alternating and synchronous tones and complexes. Thus, properties of the streaming precepts associated with these stimuli are already well-established and have been studied extensively, as we shall point out. Furthermore, the objective segregation measures we employ closely follow widely-used “deviant-detection” paradigms (***Moore and Gockel, 2002***; ***Elhilali et al., 2009***; ***Micheyl and Oxenham, 2010***; ***Carlyon et al., 2010***).

### Experiment 1: Binding the harmonic components of complex streams

In this experiment, we manipulated the streaming percepts evoked by alternating harmonic complexes of different pitches (***Singh, 1987***; ***Bregman et al., 1990***; ***Grimault et al., 2001***). We specifically investigated how attention to one of the two streams modulates the neural response to the complexes’ individual constituent tones. For example, consider the two alternating sequences of harmonic complex tones in ***Figure 2***A. The complexes in the two streams had fundamental frequencies of *F_A_ =* 400 Hz and *F_B_ =* 600 Hz, 90 ms in duration, and were separated by 20-ms gaps, with 10-ms raised-cosine onset and offset ramp. When attending to one stream, the EEG responses are known to become enhanced to the attended complexes (***Xiang et al., 2010***; ***Power et al., 2012***; ***Choi et al., 2013***). However, it is unclear whether this enhancement is due to enhanced responses to the individual tones within the attended complex or just an enhancement of the channels selectively responding to the complexes’ pitch. Conceptually, we shall hypothesize that the attentional focus on one stream effectively confers a *steady* enhancement of the responses in the frequency channels of the constituent tones, specifically those tones that are unique to the attended stream. In the next experiment, we explore the fate of tones that are shared between the two streams.

**Figure 2.**
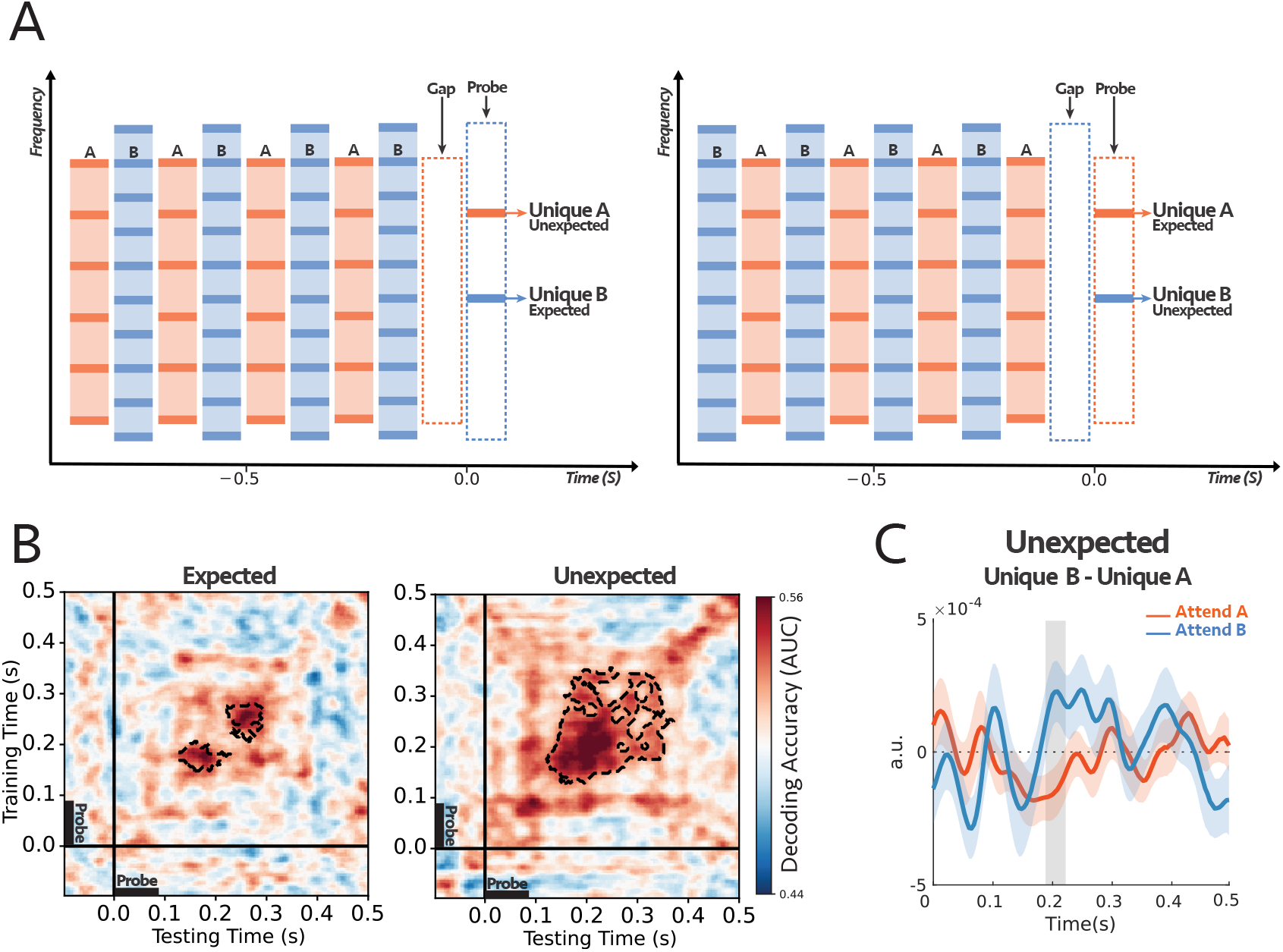
**Experiment 1: (A)** The stimulus consisted of two complex sequences with harmonically-related tones. In each trial, subjects were instructed to selectively attend to high or low pitch sequence and counted intensity deviants in the target stream. One of the frequency channels unique to tone complex A (unique A) or complex B (unique B) was probed at the end of each trial. **(B)** Decoding performance for Expected (left) and Unexpected (right) probe-tones (see the text for further explanation on the difference between the two paradigms). For each subject, classifiers were trained on the signals from all 64 EEG sensors and tested separately at each time in a 600 ms time window encompassing the probe tone (−100ms to 500ms). The trained classifiers tried to decode the target of attention within the mentioned time window. The cluster-corrected significance was contoured with a dashed line, *p =* 0.038 for expected and *p =* 0.004 for unexpected. Thus, both probes demonstrated significant attentional modulations of individual harmonic components, although the classification patterns were different between the two conditions. **(C)** Comparing the difference in responses to unique A and unique B probe-tones for two attentional conditions for the first DSS component (***Figure 2–Figure Supplement 3***; see text and Methods ***section*** for details). The comparison was significant for unexpected probes at around 200 ms after the probe onset (shaded area larger than 2*0*), with reverse polarity for *attend A* and *attend B*, suggesting the opposite effect of attention on coherent and incoherent tones (i.e., enhancement *versus* suppression). (**n=18**).

Because of the poor spatial resolution of the EEG recordings, it was difficult to investigate the attentional effects on individual frequency components during the simultaneous streams. Instead, we probed the *persistent* modulatory effects of attention on individual frequency channels immediately following the end of the streams. This was done by presenting a 90 ms pure probe-tone after a 100 ms silent gap. The probe tone was aligned with the frequency of a harmonic that is either unique to complex A (3000 Hz) or tone complex B (2000 Hz). There were two conditions for the timing of the probe-tone (***Figure 2***A): “expected”, in which the probe was a component of the *last* complex tone in the sequence (note that there is a gap between the last complex in the sequence and the probe-tone) or “unexpected” where it was a component of the *penultimate* complex tone in the sequence. The reason for these two conditions was to ascertain that the modulation of the probe-tone responses was not related to its violation of expectations (akin to the effects of “mismatched negativity“) but rather to the persistent effects of attending to one stream versus another.

18 normal-hearing participants were instructed on each trial to selectively attend to tone complex A or B and report the number of intensity deviants in the attended stream – with the deviant tone-complex is 6 dB louder than other tones in the sequence. Subjects reported hearing the two streams and being able to attend reliably to one as the behavioral results indicate, with all subjects reported the correct number of deviants above the chance level (***Figure 2–Figure Supplement 1***). To dissect the attentional effects on the responses to the two streams, we trained a set of independent logistic regression classifiers using data from all EEG sensors as explained in detail in Methods ***section*** (***King and Dehaene, 2014***; ***Stokes et al., 2015***; ***Wolff et al., 2017***). These classifiers trained on the EEG responses to the probe-tones at each time point t and tested at time t’ – where t and t’ were within the probe-tone time window (−100 ms to 500 ms) – in order to predict the target of attention. In other words, the trained classifiers tried to linearly separate the attentional conditions based on the differences within the probe topomaps. At the subject level, the classifier scores reflected the robustness of the effect across the trials (see the example in ***Figure 2–Figure Supplement 2***), and at the second level, we checked the consistency of the effect size across all subjects (***Figure 2***B). It should be noted again here that, because of the complex spatial spread of the EEG, the decoders can detect if response patterns across the scalp are modulated by the attentional focus on the unique components, but they cannot readily indicate whether the effects are simple response enhancements. For this kind of additional information, we resorted to a Denoising Source Separation (DSS) procedure to extract and examine the principle response component as detailed later below. The performance of the decoders is depicted in ***Figure 2***B. The scores of the classifiers are significantly above the chance level for both *Expected* (*p =* 0.038) and *Unexpected* (*p =* 0.004), starting from about 150 ms to 350 ms after the probe-tone onset. This performance level (up to 0.60) is commensurate with that reported in previous studies (***King et al., 2016***; ***Pinheiro-Chagas et al., 2018***). Thus, regardless of probe-tone timing and its different response dynamics due to its (expected or unexpected) context, the results demonstrate the persistent, significant differential modulatory effects of attention on the *unique* individual harmonic components of the attended and unattended sequences. We should note that the decoder significant regions differ between the two conditions of “expected” and “unexpected” probes, likely because of the differences in the detailed response patterns to the probes, as well as the effects of EEG noise which may render insignificant the response modulations at different epochs following the onset. Nevertheless, in both cases of the expected and unexpected probes, there were significant attentional modulations of the responses.

Finally, we attempted to extract an additional comparison among the probe responses under the different attentional conditions using a denoising procedure on the EEG recordings. Specifically, we isolated the most repeatable auditory component from the EEG responses to the probe-tones across trials using the DSS spatial filter (see Methods ***section***; ***De Cheveigné and Parra (2014))*** and compared the average waveform of the first DSS component and its amplitude over subjects under different conditions (see ***Figure 2–Figure Supplement 3*** for the waveforms). Importantly, we compared the *difference* in responses to the probes: *UniqueB* – *UniqueA*, under the attend A and attend B conditions. We hypothesized that, although the DSS waveform is a complex mixture of the EEG responses on all electrodes, if attention enhances coherent and suppresses incoherent responses, then the difference between the probes’ DSS responses would be modulated by attention in opposite directions, i.e., the difference: *UniqueB* – *UniqueA* would have the opposite signs for attend A and attend B, reflecting the enhancement and suppression due to attention. This was indeed the case as seen in ***Figure 2***C for the unexpected case, where the difference in probes’ responses was significantly modulated with a *reversed* polarity, at around 200 ms following the probe’s onset (shaded interval). It should be noted that the extracted responses used for the DSS differences (***Figure 2***C) and the classifiers (***Figure 2***B) are quite different, and hence it is not surprising that the detailed timing of the significance epochs following the probe-tone onsets would differ.

In the next experiment, we explore these modulations in components shared by both A and B complexes, and whether harmonicity is necessary to induce these differential attentional effects and hence play a role in segregation.

### Experiment 2: Binding of inharmonic components in complex streams

Here we extended the results of the previous experiment in several directions. *First*, we examined whether the modulatory effects of attention on the individual components of a tone complex depended on the harmonicity of the complex. *Second*, we monitored whether components *shared* between the two complex sequences experience any differential modulation. This is an important question because we had hypothesized that attention is a slow or steady-state enhancement of the components of one sequence. A shared frequency channel (by definition) belongs to both the attended and unattended streams, and hence if it is subjected to attentional effects, it must experience rapidly alternating enhancement and suppression, which would violate our hypothesis. Instead, our hypothesis predicts that shared components would not be differentially affected by selective attention to either stream. *Third*, temporal coherence is independent of the exact temporal rates or regularity of the sequences (as long as they are roughly between 2-20 Hz (***Shamma and Elhilali, 2020***). Consequently, we expected temporal coherence to be equally effective for streams of different rates (tone complexes that are temporally incoherent with each other), regardless of whether the tone complex is harmonic or inharmonic. It is, of course, expected that the streaming percept is modulated by all these parameters, i.e., whether the components of a sound are intrinsically more glued together by harmonicity regardless of streaming. By using unequal sequence rates, it was possible (as we shall elaborate) to eliminate any difference in timing expectations between the attention conditions and hence confirm the validity of the earlier results concerning the probe tones expectations.

Normal-hearing adults (21) selectively attended to one of two streams – each 90 ms in duration, with 150 ms and 250 ms inter-stimulus interval for complex A and B – based on the priming phase at the beginning of each trial, and reported whether they detected a deviant in the attended stream, ignoring deviants in the unattended stream. The two streams clearly differed by their timbre, and it was relatively easy for the listeners to track one or the other stream (see behavioral results ***Figure 3–Figure Supplement 1***). The streams ran at different rates but converged to be synchronous on the last tone in the sequence. We replaced the last tone with a single frequency probe-tone to always occur at the expected time, regardless of which stream was the target of attention (***Figure 3***A). We measured the neural responses to the probe as a function of whether its frequency belonged to one (unique) or both complexes (shared) (see ***Figure 3***A). We should note that this paradigm was effectively used previously to explore the effects of streaming on detecting timing misalignments between streams (***Elhilali et al., 2009***).

**Figure 3.**
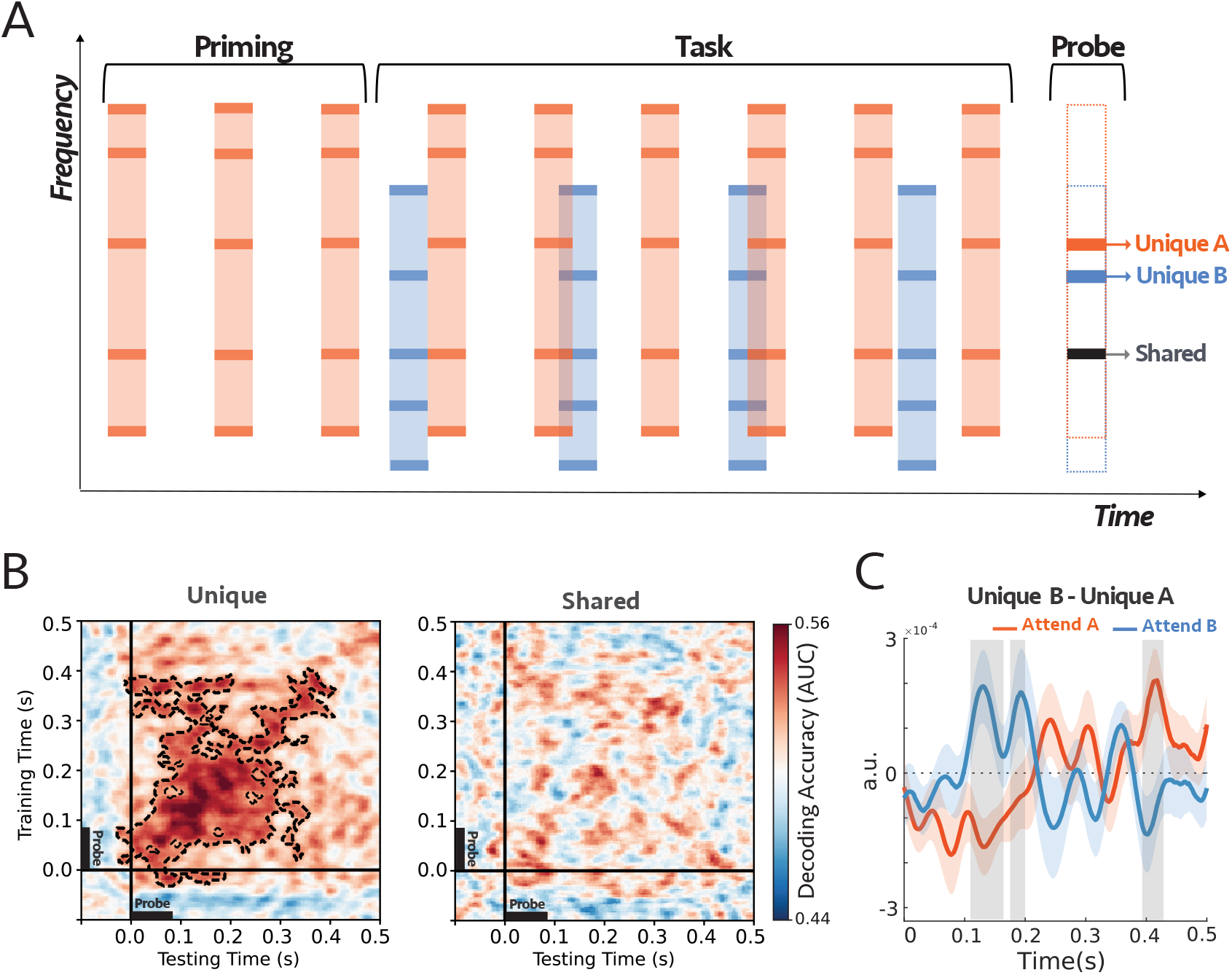
**Experiment 2: (A)** The stimuli started with a target sequence alone (Priming epoch), and after three bursts, the distractor stream was added. Both tone complexes consisted of inharmonic frequencies (see Methods ***section***). The two sequences were presented at different rates but converging on the last tone in the sequence, which was replaced by a single frequency probe-tone centered at the frequency that was either shared between the two complexes, unique to complex A (Orange), or unique to complex B (Blue). In each trial, subjects were instructed to selectively attend and detect an intensity deviant in the target sequence. **(B)** Decoding performance for unique (left) and shared (right) probe-tones. Classifiers trained and tested separately at each time in a 600 ms time window of the probe-tone (−100ms to 500ms). Cluster-corrected significance is contoured with a dashed line. The classifiers could decode attention only when the probes were unique components (*p =* 0.0003). The two sets of scores were statistically different as depicted in ***Figure 3–Figure Supplement 2***. **(C)** Comparing the difference in the first DSS component of the responses to unique A and unique B probe-tones in the two attentional conditions (***Figure 3–Figure Supplement 4***; see Methods ***section***). The comparison was highly significant for shaded areas (larger than 2*0*), with reverse polarity for *attend A* and *attend B*, suggesting the opposite effects of attention on coherent and incoherent tones (i.e., enhancement *versus* suppression). **(n=21)**.

Similar to the previous experiment’s analysis, a set of linear estimators were trained to determine the effect of attention encompassing the period of probe-tone (−100 ms to 500 ms; ***Figure 3***B). The classifiers were trained on the EEG signals to predict the attentional conditions for probe-tones, therefore for each subject, the scores summarized the effect of attention within the probe-tones (see ***Figure 3–Figure Supplement 3***). For the unique probe-tones, the performance of the classifiers in decoding attention was reliably above the chance level at 50 ms and lasted until about 400 ms after the onset of the probe, with a peak around 120 ms (*p =* 0.0003), indicating the persistent effects of attention on the unique components. However, the classifiers could not distinguish between the attention conditions when the probe-tone was shared between the two tone-complexes, suggesting that the shared frequency channels remained on average undifferentiated by the selective attention.

It is important to note that all comparisons above are between responses to the *identical probes* under different attentional conditions. Hence, we have avoided comparing the effects between the responses to the unique *versus* shared probe. However, the results of such a direct comparison are shown in ***Figure 3–Figure Supplement 2***, and they confirm the significant modulatory effects of attention on the unique probe responses relative to the shared probes.

We further analyzed the EEG responses to isolate the most repeatable neural sources with a DSS spatial filter (***Figure 3–Figure Supplement 4***A). As in the previous experiment, we compared “*UniqueB* – *UniqueA*” in attend A *versus* attend B conditions. We found the difference in responses evoked by unique A and unique B has the opposite polarity for two attentional conditions, and there are significant differences between them (***Figure 3***C). Furthermore, the average strength of the EEG response to the unique probes from 60 ms to 200 ms was significantly larger when the probe was a unique component of the attended sequence (*p =* 0.03 for Unique A and *p =* 0.01 for Unique B), but no significant difference was observed in response to shared channels when the subject attended to tone complex A or tone complex B (*p =* 0.6; ***Figure 3–Figure Supplement 4***B). These results, therefore, confirm that attention relatively modulates the unique frequency components of the attended complex. We emphasize that this attentional modulation occurred despite the *inharmonicity* of the components, and hence we conclude that these effects are due to the temporal coherence of the tone components and not any harmonic relation among them. In addition, consistent with the temporal coherence predictions made above, the shared component remained undifferentiated by the attentional focus.

### Experiment 3: Binding Noise Sequences with coherent tones sequences

Experiments 1 and 2 confirmed that the binding of the components within a stream relies primarily on their temporal coherence and not on any harmonic relationship among them and that different sequences segregate well when running at different rates. Here, we investigate binding one step further to demonstrate that temporally coherent sound elements bind perceptually to form a stream even when they are of a different nature, e.g., tones and noise-bursts, and even when placed far-apart along the tonotopic axis. Specifically, Experiment 3 tested the hypothesis that attending to a distinct stream of noise bursts not only will modulate its neural responses it also affects all others that are temporally coherent with it, effectively binding the percept of the noise with the coherent tones to form a single unified stream. Moreover, we tested again if shared tones remain uncommitted as we found earlier, hence contributing equally to both streams.

***Figure 4***A illustrates the stimuli and procedures used with 14 normal-hearing subjects who were instructed to focus attention on a sequence of narrow-band noise bursts – between 2-3 kHz, 125 ms tone duration, and 250 ms inter-stimulus interval – and report the number of deviants that occurred in the noise stream. Two sequences of inharmonic tone complexes (A and B) accompanied the noise sequence, one coherent with the noise (the attended stream) and one alternating (incoherent) with it. At the end of each trial, a single frequency probe-tone was presented at frequencies aligned with either a unique component of complexes A or B or a component shared by both complexes. All subjects perceived the streaming of the alternating stimuli and performed the task above the chance level as demonstrated by the behavioral results in ***Figure 4–Figure Supplement 1***.

**Figure 4.**
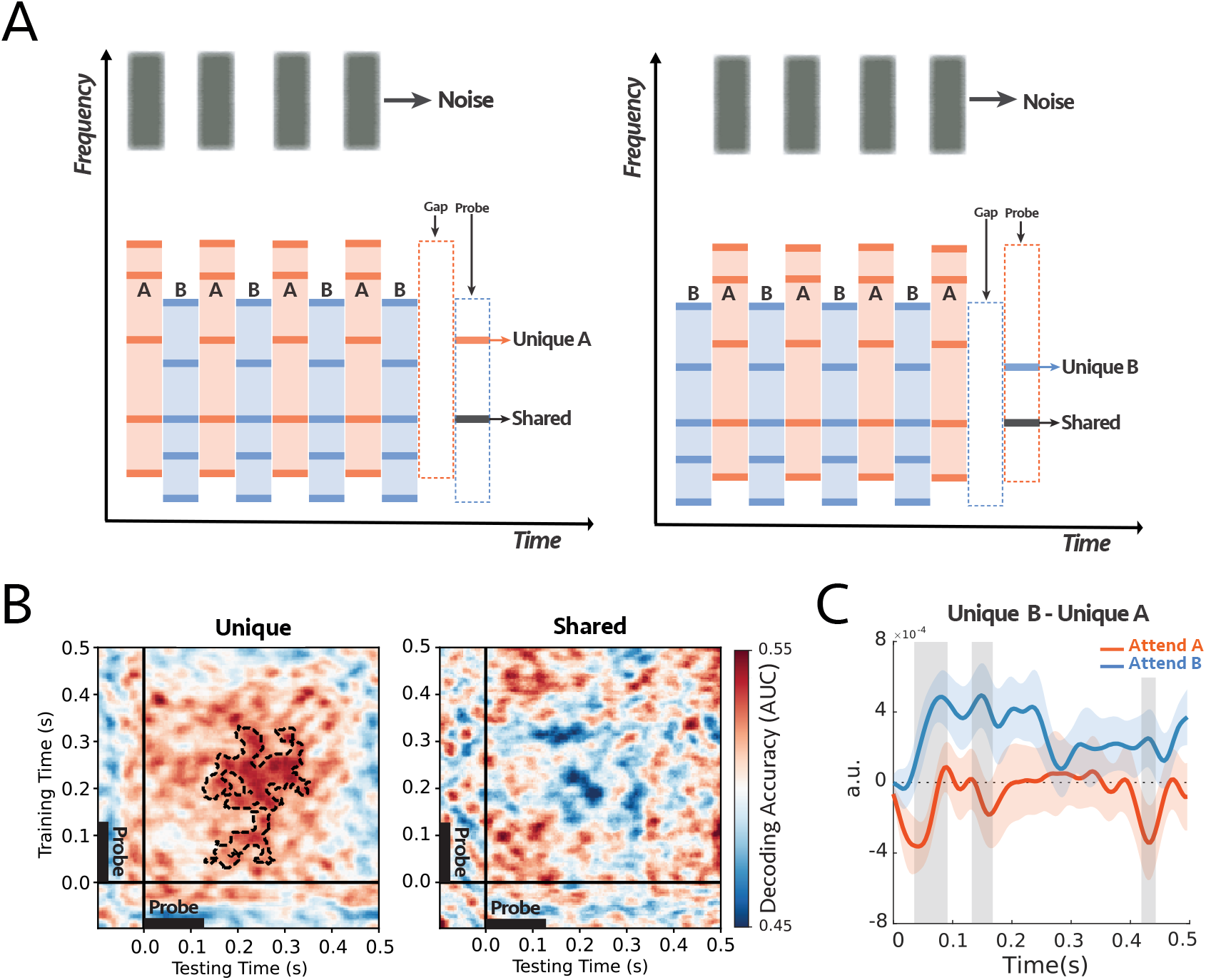
**Experiment 3: (A)** The stimulus consisted of two complex sequences with inharmonic frequencies - the frequency components of the complex tones were the same as experiment 2 - and a sequence of noise as a target stream. In each trial, subjects were instructed to always attend and count intensity deviants in the target noise sequence. At the end of each trial, we probed a unique frequency channel to tone complex A or complex B or shared between the two complex tones. **(B)** Decoding performance for unique (left) and shared (right) probe-tones. Classifiers trained and tested separately at each time in a 600 ms time window of the probe-tone (−100ms to 500ms). The significant cluster is contoured with a dashed line. The classifiers could decode attention only when the probes are unique components (*p =* 0.004). There is a statistical difference between the unique and shared scores as depicted in ***Figure 4–Figure Supplement 2***. **(C)** Comparison between the difference in responses to unique A and unique B probe-tones for two attentional conditions for the first DSS component (***Figure 4–Figure Supplement 3***; see Methods ***section***). There were significant differences for shaded areas (larger than 2*0*), with reverse polarity for *attend A* and *attend B*, suggesting the opposite effect of attention on coherent and incoherent tones (i.e., enhancement and suppression). **(n=14)**.

As before, we trained classifiers to detect modulations on the probe tone responses during, pre, and post the probe tone onsets (at 0 ms). Results displayed in (***Figure 4***B) demonstrate that attention significantly modulates the probe-tone responses starting 130 ms following onset but only when aligned with unique tones of the complex-tone sequences (*p =* 0.004; ***Figure 4***B, left panel). The classifier failed to decode any modulations due to attention when the probe-tone aligned with a shared component (***Figure 4***B, right panel). Moreover, there was a statistical difference between the decoder scores of Unique and Shared probe tones, as illustrated in ***Figure 4–Figure Supplement 2***.

We then analyzed the EEG responses to extract the most repeatable auditory component across trials by looking at the first DSS component (see Methods ***section***), which was averaged across all subjects (***Figure 4–Figure Supplement 3***A). ***Figure 4***C illustrates the difference between the unique probes *UniqueB* – *UniqueA* under attend A (orange) and attend B(blue) conditions. For most of the time period encompassing the probe tone, the difference exhibited the opposite polarity in the two attention conditions, with high significance near 70 ms, 160 ms, and 420 ms following the probe’s onset. Furthermore, we looked at the *average* of the first DSS components over the time window of 60 ms to 200 ms. The average power is significantly larger when the probe-tone is a unique component of the attended stream (*p =* 0.04 for unique A and *p =* 0.01 for unique B). However, there is no significant difference in response to the shared frequency channel (*p =* 0.24; ***Figure 4–Figure Supplement 3***B).

To summarize, the key finding of this experiment is that attending to the noise-bursts, which are perceptually different from the tones and spectrally located at least 2.5 octaves apart, nevertheless caused the coherent complex-tone sequences to become modulated as if they became bound to the noise-bursts and included in the focus of attention. This is consistent with the earlier experiments’ findings in the present study that coherent tones are modulated when subjects attended directly to the complexes. This we take as evidence of the perceptual binding of all coherent acoustic components to form a unified attended stream.

### Experiment 4: Segregating Speech Mixtures

Real-world auditory scenes often consist of sound streams of unequal rates, many shared spectral components, and gradually changing parameters (pitch, location, or timbre). In all previous experiments, we have demonstrated that temporal coherence plays a crucial role in stream formation. However, all stimuli used were well-controlled, relatively simple tone-complexes and noise bursts with stationary parameters. Here, we extend the temporal coherence principle tests to a more naturalistic context using speech mixtures. In a speech, the signal is modulated in power during the succession of syllables, just like the tone and noise sequences used in the previous experiments, i.e., one can abstractly view a speech signal as a sequence of bursts separated by gaps of various durations. Each burst encodes features of the speaker’s voice, such as his/her pitch, location, and timbre, which temporally fluctuate in power coherently. In a mixture of two different speakers, female (F) and male (M), saying different words, the sequences of bursts begin to resemble the alternating A & B complexes of our simpler stimuli. Consequently, the power in each speaker’s features would fluctuate coherently, but they are both different and out-of-sync with those of the other speaker. Furthermore, simultaneous speech segregation can potentially rely on the incoherence between the power modulations of the two speech streams since speakers utter different words, and hence their modulations are often de-synchronized (***Krishnan et al., 2014***).

To confirm these assertions, we first tested that the same approaches using probe-tones and trained classifiers can be readily applied to decoding speech responses. The probe-tones were restricted to the end of the sentences and were always harmonic complexes, as detailed below. In the second part of experiment 4, we refined the probe-tones to investigate the attentional modulations on single frequency components, and more importantly, to insert the probe anywhere in the midst of the speech mixture.

#### a) Probe at the end

Single-speaker sentences were selected from the CRM corpus (***Bolia et al., 2000***) and then mixed to produce two-speaker mixtures, each containing a male and a female voice. All sentences in this corpus have the same format, including color and a number (see Methods ***section***). During the task, subjects were instructed to attend to a specific speaker on each trial and then report the color and number uttered by this target speaker. The mixture in each trial ended with a 90 ms harmonic-complex tone as a probe, consisting of the 4 lowest harmonics aligned with corresponding frequency components of either the attended or unattended voice. Therefore, the probe-tone was uniquely aligned with one speaker, as were the single-tone probes in the earlier experiments (***Figure 5***A).

**Figure 5.**
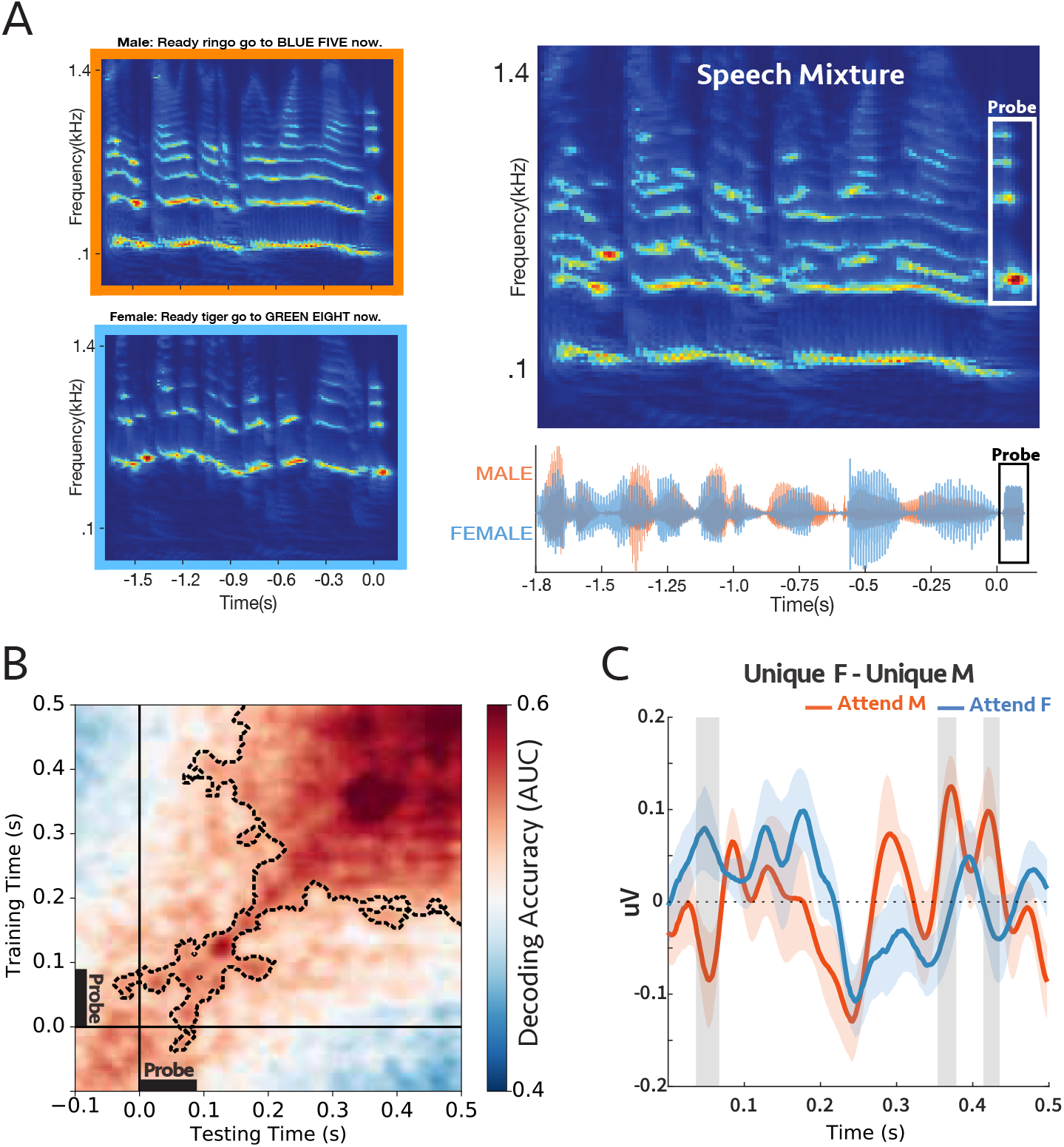
**Experiment 4a (A)** A sample stimulus for experiment 4a. *Top-left*: cochlear spectrogram of the male voice. *Bottom-left*: cochlear spectrogram of the female voice. Note the probe-tone at the end. *Right*: cochlear spectrogram and acoustic waveform of the mixture. All the sentences had the same length (1.8 sec). The participants were instructed to report the color or the number by the target voice. During the task, the auditory scene consisted of two concurrent speech streams followed by a 90 ms probe-tone with complex harmonics. The probe-tones harmonic frequencies were aligned with the four loudest harmonic frequencies of either male or female voice at the end of the sentences; therefore, the probe-tone was either unique to male or unique to female. **(B)** Decoding performance for the probe-tones trained and tested during the probe time window (−100ms to 500ms). The significant clusters were contoured with a dashed line. The classifiers could decode attention significantly above chance (*p =* 0.0002) **(C)** Comparison between the difference in evoked responses to unique F and unique M probe-tones for two attentional conditions at *Cz* channel (placed on the center of the mid-line sagittal plane) (***Figure 5–Figure Supplement 3***; see Methods ***section***). There were significant differences for shaded areas (larger than 2*0*), suggesting the opposite effect of attention on coherent and incoherent tones. **(n=16)**.

16 Participants were asked to report the color and the number mentioned in the target sentence to make sure that they were attending correctly to the target speaker, all the subjects were able to do the task with ease (average accuracy = %93; ***Figure 5–Figure Supplement 1***). Meanwhile, we measured the neural responses to the probe-tone with EEG. The responses were compared under different attention conditions using the same linear classifiers described earlier, with decoder scores significantly above the chance level (***Figure 5***B). Additionally, we generalized the modulatory pattern of attention by using classifiers trained and tested at various times relative to the probeonset, e.g., trained at the beginning of the speech mixture (−1.8 to −1.2 s) and tested around the probes (−0.1 to 0.5s; ***Figure 5–Figure Supplement 2*** left panel), trained near the probe (−0.1 to 0.5s) and tested at the onset of the speech mixture (−1.8 to −1.2 s; ***Figure 5–Figure Supplement 2*** right panel) In all cases above, the decoding scores were significantly above chance as is evident in the figures (*p =* 0.0002, *p =* 0.009, and *p =* 0.009 respectively). We also contrasted the evoked responses to the probe-tones (*UniqueF* – *UniqueM*) at the Cz channel and observed significant differences between attending to male and female, with opposite polarity (***Figure 5***C).

Therefore, the results thus far indicate that: 1) The neural response to the harmonic frequency components aligned to the pitch of the two speakers is reliably modulated by attention. 2) The pattern of brain activity at the onset of the attended/unattended speech sentence is similar to the activity during the probe tone at the end, and consequently, the trained decoders were generalizable for these two time windows even when separated by a sizable interval. These results are consistent with the temporal coherence hypothesis because if attention to the pitch of one voice enhances the pitch signal, it will enhance its harmonics (all being coherent with it; ***Krishnan et al.*** (***2014***)) and will relatively suppress the harmonics of the unattended speaker, which are incoherent with it.

#### b) Probe in the middle

Using the same speech corpus as stimuli, this experiment probed the modulations of a single frequency channel potentially anywhere within the duration of the speech mixture. The probe frequencies in these experiments were chosen centered at the 2nd harmonic of the female or at the 3rd harmonic of the male, unique components in the midst of the speech mixtures as illustrated in ***Figure 6***A. Participants were instructed to report the color or number spoken by the talker who uttered the target call-sign (“Ringo”; see Methods ***section***), all participants did the task successfully (average accuracy = %79; ***Figure 6–Figure Supplement 1***). On average, the onset of the call-sign occurred 300 ms (± 25 ms) following sentence onset, and the probe-tone was inserted 600 ms after speech onset.

**Figure 6.**
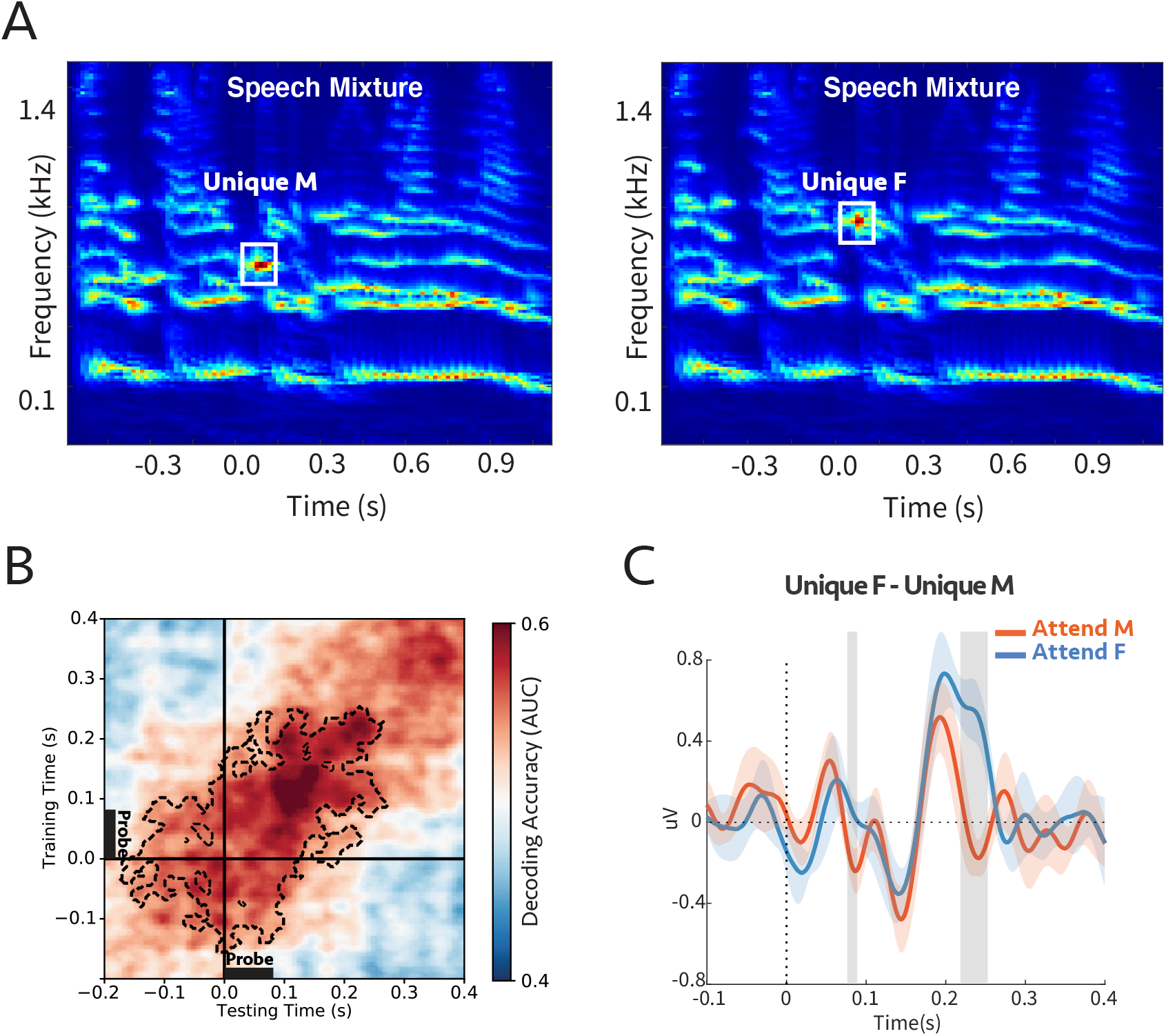
**Experiment 4b (A)** Cochlear spectrogram of a sample stimulus mixture for experiment 4b. It consisted of two (male and female) voices. The participants were instructed to report the color or the number of the speaker who uttered the target call-sign. The probe’s frequency was aligned with the 2nd or 3rd harmonic of the female or male, respectively. Therefore the probe-tone was either unique to male (unique M) or unique to female (unique F). **(B)** Decoding performance for the probe-tones. Classifiers trained and tested separately at each time in a 600 ms time window of the probe-tone (−200ms to 400ms). Cluster-corrected significance was contoured with a dashed line. The classifiers could decode attention significantly above chance for up to 280 ms after the probe-tone onset (*p =* 0.015). **(C)** Comparison between the difference in evoked responses to unique F and unique M probe-tones for two attentional conditions at *Cz* channel (placed on the center of the mid-line sagittal plane) (***Figure 6–Figure Supplement 2***; see Methods ***section***). There are significant differences for shaded areas (larger than 2*0*), suggesting the opposite effect of attention on coherent and incoherent tones. **(n=7)**.

We trained linear classifiers to ascertain the modulation that attention induced in the probe responses during the time window (−200 ms to +400 ms with probe onset defined as 0). It is evident in ***Figure 6***B) that the decoding scores were significantly above the chance level and lasted for up to 280 ms with a peak at 150 ms after the probe-tone onset. We also extracted the evoked EEG signal at channel Cz due to the probe-tones to determine the direction of the modulation induced by attention ***Figure 6–Figure Supplement 2***. ***Figure 6***C shows the difference in response to unique F and unique M was significantly and rapidly modulated by attention within about 250 ms, consistent with previous findings in ECoG recordings (***Mesgarani and Chang, 2012***).

Therefore, in conclusion, we measured significant attentional modulations of the probe-tone responses that are frequency-specific (distinguishing between alignments with closely-spaced male and female harmonics). These findings indicate that during speech segregation, the components of the attended speaker are differentiated from those of the unattended source quite rapidly, or specifically, as soon as 250 ms after the onset of the target callsign. This delay is commensurate with that observed in analogous ECoG experiments involving switching attention between 2 speakers.

## Discussion

This study explored the dynamics and role of temporal coherence and attention in the binding of features that belong to one auditory source and its segregation from background sources in a cluttered auditory scene. The temporal coherence hypothesis predicts that acoustic features that induce coherently-modulated neural responses should bind together perceptually to form a single stream. One piece of evidence for this perceptual stream formation is taken to be the physiological enhancement of all the coherent responses. It is also postulated that an essential component of this process is attention directed to one or more of the coherent set of acoustic features, which then initiate mutual interactions, and hence binding of all coherent features. Previous studies have shown that responses to an attended (pure-tone) stream are enhanced relative to the remaining background in a mixture (***Ding and Simon, 2012***; ***Snyder et al., 2006***). However, it has remained unclear whether the neural responses to the individual constituents of a complex stream are also similarly modulated by attention to only one of its elements and how this is related to its perceptual formation.

In the series of EEG experiments reported here, we demonstrated that when listeners attended to one attribute of a complex sound sequence, other temporally coherent responses were similarly modulated; incoherent responses were relatively suppressed (or *oppositely* modulated) while leaving shared elements unchanged. This was found to be true over a wide range of attributes, be it the pitch of a sequence of harmonic complexes (Experiment 1), the timbre of inharmonic complexes (Experiment 2), noise burst sequence (Experiment 3), or the call-sign by a single speaker in a mixture (Experiment 4). Of crucial importance, this was the case even for those features in the sound sequence that were *not* directly accessible to the listener. For example, when subjects attended to the pitch of a harmonic complex or the timbre of an inharmonic complex, they rarely reported being able to (or spontaneously) listen to the individual constituent tones, yet these components became modulated as if they were directly attended to. In fact, in experiment 3, subjects completely ignored the accompanying complex tones while attending only to the noise bursts, yet response modulations still occurred for the unique coherent tones, i.e., they acted as part of the foreground noise stream.

To access the responses of the individual components of a stream (despite the poor spatial resolution of the EEG), we investigated the responses to probe tones and probe complexes that relied on the persistent effects of attention in the midst or just following the end of the streams when there were no interfering signals from other sounds. The effects of attention on the probe responses, however, were not always easy to interpret because the array of the 64 EEG electrodes pick up complex mixed signals deriving from many regions of the brain. Thus, specifying and interpreting an EEG response to use for the measurement requires combining (and not simply averaging) the recordings from all these electrodes. Therefore, the term “response enhancements” used in our original hypothesis does not always literally mean an *increased* response amplitude or power, but rather a response-modification that is robust and repeatable when attentional conditions are identically manipulated. While these changes are often detected as enhancements in the power of the response DSS component (particularly when using simple pure-tone streams (***Xiang et al., 2010***; ***Power et al., 2012***) instead of the complex multi-component streams here), we focused instead on a more flexible measurement approach that detects these changes through linear estimators. Specifically, a set of classifiers were trained to decode the attended/unattended responses near the probe-tone time window, and were then tested at other times (such as generalizing the estimators to the speech beginning) to demonstrate that the response patterns induced by attention persisted during and subsequent to the probe.

To summarize, the overall findings from this study are consistent with the temporal coherence hypothesis where correlated responses become bound as a single stream that integrates the elements of the sequences regardless of: (1) the temporal regularity of the sequences, e.g., uniform or irregular (***Figure 2*** and ***Figure 3***); (2) stimulus types across the sequences, e.g., noise or tones-complexes (***Figure 4***); (3) whether the tones are harmonic or inharmonic complexes (***Figure 2***, ***Figure 3***, and ***Figure 4***); (4) whether the sequences are spectrally near or far apart (***Figure 2***, ***Figure 3***, ***Figure 4***, and ***Figure 5***); and crucially, (5) whether the sequence parameters (e.g., pitch and timbre) are stationary or dynamically slowly evolving as in speech (***Figure 5*** and ***Figure 6***).

Temporal coherence is essentially an associative process likely enabled by rapidly formed and modulated connectivity among coherently responsive neurons. This process is analogous to the well-known Hebb’s rule of “*fire together, wire together*”, except that it occurs at a much faster pace (within hundreds of milliseconds, as evidenced by the rapid build-up following the call-signs in ***Figure 6***). It is also promoted and controlled by “attention”, a notion that is difficult to define precisely. However, experiments in animals and human subjects have demonstrated that without the engagement and attentional focus on the task, or selective attention to specific features of the stimuli, these rapid modulations of connectivity which are manifested as perceptual binding, and hence stream formation, become far weaker or absent (***O’Sullivan et al., 2015***; ***Lu et al., 2017***). The underlying biological foundations of this process remain largely unknown but are currently the target of numerous ongoing studies.

We end by noting that the concept of temporal coherence likely applies in a similar way in other sensory modalities such as vision. In a dynamic visual scene, features of a visual object, such as its pixels, move together coherently in the same direction and speed, inducing highly correlated neural responses. Conversely, pixels of independent objects move with different relative motion and can thus be segregated easily from those of other objects based on this idea (***Lee and Blake, 1999***). Also, multi-modal integration, such as enhanced comprehension of speech in an audio-visual scenario (lip-reading), may well be explained by temporal coherence, i.e., the temporal coincidence between the visual representation of the lip motion and the acoustic features of the syllables can strongly bind these sensory features, and hence improve the intelligibility of speech (***Bernstein et al., 2004***; ***Crosse et al., 2016***; ***Atilgan et al., 2018***; ***O’Sullivan et al., 2020***).

## Methods

### Terminology

In the present study, several terms are used somewhat interchangeably to refer to the idea of temporal coherence, which more specifically states that neural responses that are temporally correlated over a short period of time on a pair of sensory pathways can evoke a unified percept or “become bound” into a single stream. Sometimes the reference is made instead to the stimuli that evoke these responses, and hence terms like synchronized tone sequences or coincident events occurring over a period of time can mean the same thing as temporal coherence. In all these cases, the context will hopefully clarify the intent as it is by no means necessary that any of these stimuli can unambiguously evoke the necessary correlated activity. For instance, a single pair of synchronized events are irrelevant to stream formation since coincidence must occur multiple times over a short interval. Similarly, synchronized bursts of random tone complexes do not evoke coherently modulated activity on any pair of frequency channels and hence do not bind. For more examples of such conditions, please see ***Shamma et al. (2011)***.

### Participants

76 young adults with normal-hearing (ages between 19 and 31) participated in this study, consisting of four experiments. 18, 14, 21, and 23 subjects participated in experiments 1-4, respectively. Experiments 1 and 3 were conducted at the University of Minnesota and experiments 2 and 4 were conducted at the University of Maryland. All participants were given course credits or monetary compensation for their participation. The experimental procedures were approved by the University of Maryland and the University of Minnesota Institutional Review boards. Written, informed consent was obtained from each subject before the experiment.

### Data Acquisition and Stimuli Presentation

Data were collected at two sites. At University of Maryland, Electroencephalogram (EEG) data were recorded using a 64-channel system (ActiCap, BrainProducts) at a sampling rate of 500 Hz with one ground electrode and referenced to the average. We used a default fabric head-cap that holds the electrodes (EasyCap, Equidistant layout).

EEG data from University of Minnesota were recorded from 64 scalp electrodes in an elastic cap, using a BioSemi ActiveTwo (BioSemi Instrumentation). EEG signals were acquired at a sampling rate of 512 Hz, and referenced to the average. We analyzed the EEG data offline.

The stimuli were designed in MATLAB and presented to the participants with the Psyctoolbox(***Brainard, 1997***; ***Pelli, 1997***; ***Kleiner et al., 2007***). The stimuli audio was delivered to the subjects via Etymotics Research ER-2 insert earphones at a comfortable loudness level (70dB).

### Stimuli Design

#### Experiment 1

The stimulus was presented as an alternating ABAB sequence with a sampling rate of 24414 Hz, followed by a short probe-tone. Segment **A** was a harmonic tone complex with a fundamental frequency (F0) of 400 Hz, while **B** was a harmonic tone complex with an F0 of 600 Hz. All A and B harmonic complexes were generated with random starting phases and were low-pass filtered to exclude frequency components higher than 4000 Hz with a 48 dB/oct filter slope. Each segment of A or B lasted 90 ms, including 10-ms raised-cosine onset and offset ramps. Segments were separated by 20-ms gaps (the repetition rate of both A and B tones alone was 4.55 Hz). The total number of AB harmonic tone pairs in one trial was randomly chosen from 27 to 33, so participants could not predict the sequence’s total duration. The sequence was followed by a 100-ms silent gap and a 90-ms pure-tone probe. The level of probe-tones and each component of A and B harmonic complexes were at a *rms* level of 55 dB SPL. The ending tone complex in the sequence was balanced so that half the trials ended with tone complex A. The other half ended with tone complex B (see ***Figure 2***A), thus based on these structures, we defined 2 different conditions for the relative position of the probe and the last complex tone in the sequence: (1) When the probe tone is a component of the last complex, and therefore it occurred at the supposedly *expected* spectral location (e.g., the sequence ended with tone complex A and we probed the frequency channel unique to complex A), and conversely (2) when the probe tone was a component of the penultimate complex tone and occurred at an *unexpected* spectral location (e.g., the sequence ended with tone complex B and we probed the frequency channel unique to complex A).

The experiment included six blocks of 100 trials each. Participants were instructed to selectively attend to the A tones (the higher pitch) in half the blocks and the B tones (the lower pitch) in the other half of the blocks by explicitly ask them to pay attention to the higher or lower pitch sequence. The order of blocks was randomized. The 300 trials for each set of instructions were equally divided into two groups, depending on the probe-tone used. The two probe-tones were 2000 Hz (unique frequency component of A) and 3000 Hz (unique component of B). The order of trials with different probe-tones was randomized within the three blocks for each set of instructions. We used intensity deviant in both A and B harmonic tones to monitor stream segregation and attention. There were 0 - 3 deviants tones in each sequence, and the number of deviants was uniformly distributed across trials. Out of 100 trials within a block, there were 25 trials with intensity deviant segments only in the A sequence, 25 trials with deviants only in the B sequence, 25 trials with deviants in both sequences, and 25 trials with no deviant. Deviant tones were presented at a level 6 dB higher than the regular tones. Participants were instructed to press a button (0, 1, 2, or 3) after hearing the probe-tone at the end of each sequence to answer how many intensity deviants they detected within the attended stream while ignoring deviants in the unattended stream. Deviants were prevented from occurring in the first five and last three tone pairs. To analyze the EEG data, we kept the trials in which participants reported correctly, e.g., the exact number of deviants in the target sequence.

#### Experiment 2

The auditory scene consisted of two complex tones denoted as A and B and presented at a sampling rate of 44100 Hz, 90 ms in duration, 10 ms cosine ramp, and 150 ms onset-to-onset interval for tone complex A (6.67 Hz repetition rate) and 250 ms interval for tone complex B (4 Hz repetition rate). Each complex tone consisted of 5 predefined inharmonic frequencies with one shared frequency component between the two complexes (*F_A_*=[150, 345, 425, 660, 840] Hz and *F_B_*=[250, 425, 775, 1025, 1175] Hz). The two sequences were presented at different rates but converged on the last tone in the sequence. A single frequency probe-tone replaced the converged tones; thus, the probe-tone always occurred at the expected location (see descriptions for experiment 1). The probe-tone was centered at the frequency that was shared between the two complexes (425 Hz) or at a frequency unique to complex A (660 Hz) or complex B (775 Hz). The participants were instructed to pay attention to a complex sequence that was included in the priming epoch and report whether they heard an intensity deviant (5 dB increase in the loudness) only in the target sequence, with a 20% chance of having a deviant in complex A sequence and independently 20% chance of having a deviant in the sequence of complex B. Deviant was prevented from occurring in the first and last two tones; to analyze the EEG data, we only kept the trials in which participants reported correctly (i.e., all trials with missed and false alarms were discarded). Each trial started with a target stream alone (priming), and after three bursts, the distractor stream was added (***Figure 3***A). The priming phase was balanced for all trials, so half the trials started with tone complex A, and the other half started with tone complex B. Trials duration were uniformly distributed between 3.5-5 sec to avoid the formation of expectation of the ending.

The experiment was conducted in six blocks of 100 trials, and attention was fixed on one stream throughout an entire block. The probe-tone frequency was uniformly selected from [425 Hz, 660 Hz, 775 Hz] for all trials. Before neural data collection, a training module was provided. Subjects received feedback after each trial for both training and the test sets.

#### Experiment 3

This experiment consisted of a narrowband noise sequence with a passband of 2-3 kHz. The noise was accompanied by two inharmonic complex sequences, with one of them coherent with the noise sequence, and the other complex was alternating. We used the same frequency components as *experiment 2* for both inharmonic complexes (*F_A_*=[150, 345, 425, 660, 840] Hz and *F_B_*=[250, 425, 775, 1025, 1175] Hz presented at a sampling rate of 24414 Hz, and a level of 70 dB). Each noise segment and tone complex was 125 ms in duration, including a 15 ms offset and onset cosine ramp with a 250 ms onset-to-onset time interval for all tones. We used 0 - 3 (with equal probability) intensity deviants, a 6dB increase in amplitude, both in the target of attention (noise) and the distractor (alternating complex). The subjects’ task was always to pay attention to the noise sequence and count the number of intensity deviants only in the attention target (noise). To analyze the EEG data, we only kept the trials where subjects reported the *exact* number of deviants. The trials’ duration was uniformly distributed between 3.5-5 secs, so participants could not form an expectation for the sequence’s total duration. We inserted a single frequency probe-tone 125 ms after the last tone complex in the sequence. The probe-tone was at the frequency shared between the two complexes (425 Hz) or was unique to complex A (660 Hz) or complex B (775 Hz). To ensure that the EEG response to the last complex tone does not affect the probe-tone response under different attention conditions, trials ended with complex tone A when the probe-tone was unique to B and ended with complex tone B when the probe-tone was unique to A.

The experiment was conducted in six blocks of 100 trials. For 3 blocks, complex A and the rest of the blocks complex B were coherent with the noise sequence. The probe-tone frequency was uniformly selected from [425 Hz, 660 Hz, 775 Hz] for all trials within a block. Before neural data collection, a training module was provided. Subjects received feedback after each trial for both training and test sets.

#### Experiment 4

We used the CRM speech corpus (***Bolia et al., 2000***) for this experiment. In general, this speech database consists of 8 different speakers (4 female), and the format of each speech sentences is: “Ready [Callsign] go to [Color] [Number] now”. The callsign set is: ‘Charlie’, ‘Ringo’, ‘Laker’, ‘Hopper’, ‘Arrow’, ‘Tiger’, ‘Eagle’, ‘Baron’. The color set is: ‘Blue’, ‘Red’, ‘White’, ‘Green’, and the number set is: {1, 2, 3, 4, 5, 6, 7, 8}. Therefore, each speaker has 256 unique sentences. Two speakers (one female and one male) were chosen for the task. We manipulated each sentence’s duration, so all the sentences had the same length (1.8 seconds at a sampling rate of 40000 Hz), and on average, the Callsign occurred 300 ± 25 ms after the speech onset.

For both parts, we only kept the trials in which participants reported correctly for the EEG analysis, i.e., if the listeners’ reports matched with the color or number uttered by the target speaker (see below).

Part a) During the task, the auditory scene consisted of two concurrent speech streams - we constrained the mixtures to have different ‘callsigns’, ‘colors’ or ‘numbers’ in male and female voices - that were followed by a probe-tone with complex harmonics. The probe tone’s harmonic frequencies were aligned with the 4 loudest harmonic frequencies of either male or female voice at the end of the sentences; therefore, the probe-tone was unique to male or unique to female voices. The probe’s duration was 90ms, with a 10 ms cosine ramp, and played after a 10ms interval right after the sentences (***Figure 5***A). The experiment was conducted in 4 blocks with 100 trials. Participants fixed their attention on either male or female voice for 100 trials in a given block, and they reported the color or the number of the attended speaker that was asked randomly at the end of each trial. The order of the blocks was shuffled across subjects.
Part b) In the second part of the experiment, we inserted a single frequency probe-tone in the middle, following 600 ms of the speech onset and around 300 ms after the callsign onset. The probe-tone was 90 ms in duration, including a 10 ms cosine ramp, followed by a 10 ms gap of silence. The frequency of the probe-tone was aligned with the 2^*nd*^ harmonic of the female voice (unique to female, average *F_F_ =* 391*Hz*) or the 3^*rd*^ harmonic of the male voice (unique to male, average *F_M_ =* 288*Hz*). On average, the callsign occurred 300*ms* ± 25*ms* after speech onset. Although the probe tone was presented in the speech, the speech mixture was masked by complete silence for the probe-tone duration. The experiment was conducted in a block of 400 trials, and participants were instructed to pay attention to the speaker who uttered the target callsign (‘Ringo’) and report either the color or number (randomly selected for each trial) spoken by the target voice, at the end of each trial.

### EEG Prepossessing

After loading, EEG data were mean-centered. The bad channels were interpolated based on the data from the neighbor channels. The slow varying trend in data was removed by robust fitting a polynomial (***de Cheveigné and Arzounian, 2018***). For the DSS analysis (see below ***subsection***), data were bandpass filtered between 1 Hz to 20 Hz with Butterworth window of order 4 using ‘filtfilt’ in MATLAB. Eyeblink components were isolated and projected out with the HEoG and VEoG channels using a time-shift PCA (***de Cheveigné and Simon, 2007***). Data were referenced by subtracting the robust mean and epoched based on the triggers sent at the beginning of each trial. Finally, the outlier trials (bad epochs) were manually detected and discarded based on a threshold criterion.

### Decoding

Decoding analysis was performed using sci-kitlearn (***Pedregosa et al., 2011***) and MNE (***Gramfort et al., 2013***) libraries in python 3.6. We trained linear classifiers on EEG sensor space signals, band passed 0.1-20 Hz at 250 Hz sampling frequency (***King and Dehaene, 2014***). At each time point *t*, we trained a classifier using the matrix of observations *X_t_* ∈ *R*^*N*×64^, for 64 electrodes in *N* samples, to predict the vector of labels *y_t′_* ∈ {0, 1}^*N*^ at every time point *t′* in a trial. The labels correspond to the two attention conditions (attend A versus attend B in experiment 1, 2, and 3 or attend female versus attend male in experiment 4). For example, for each subject, we trained the decoders on EEG signals at time points encompassing the probe tone (−100 ms – 500 ms). Therefore, the decoder at each time point learns to predict the attended stream using the EEG sensor topography at the same time point. Then, we generalized the trained decoder by testing it on all other time points of the trial. Logistic regression classifiers were used, with 5-fold cross-validation, within-subject for all the trials. We used the area under the receiver operating characteristic curve (AUC) to quantify the classifiers’ performance.

In summary, within a subject, the classifiers’ scores imply the robustness of the attentional effects on the probe-tone response topography. So the significant time regions in all figures corresponding to decoder scores indicate the effect’s consistency across all subjects.

### Denoising Source Separation (DSS)

A set of spatial filters are synthesized using a blind source separation method that allows the measurement of interest to guide the source separation. For detailed explanation see ***De Cheveigné and Simon (2008)***. For our purpose, the Denoised Source Separation (DSS) filter’s output is the weighted sum of the signals from the 64 EEG electrodes, in which the weights are optimized to extract the repeated neural activity across trials. Therefore, for the experiment 1, 2, and 3, the first DSS component reflects a brain source of auditory processing, repeatedly evoked during the segregation task for the same set of sound frequencies.

Our use of the DSS method required a large number of the same stimuli to extract the repeated activity. However, in our speech experiment (experiment 4a and 4b), since each trial consisted of various sentences with varying sound frequencies, different neural activities were driven by the stimulus in different trials; therefore, it is difficult to isolate the first DSS component as we did for tone experiments. Thereby, we only used the DSS method in order to denoise the data in experiment 4, in which we projected back the first 5 DSS components to the sensor space to form a clean and denoised dataset. Finally, we compared the evoked responses at the Cz channel (placed on the center of the mid-line sagittal plane) for experiment 4.

### Statistical Analysis

Statistical analysis for the decoders was performed with a one-sample t-test with random-effect Monte-Carlo cluster statistics for multiple comparison correction using the default parameters of the MNE spatio_temporal_cluster_1samp_test function (***Maris and Oostenveld, 2007***). To compare the differences in evoked responses due to the probe-tones, we performed bootstrap resampling to estimate the standard deviation (SD) of the difference between the attention conditions. We checked whether at each time point the difference between attention conditions exceeded 2×estimated SD (2*σ*). In supplement figures, for the first DSS component’s strength comparison between two attention conditions, we used a one-tail non-parametric Wilcoxon signed-rank test (***Wilcoxon, 1945***). Error bars in all figures are ±SEM (standard error of the mean).

## Acknowledgments

We are grateful to Andrew Oxenham and Hao Lu for their critical contributions in the earlier stages of this work, the experiments’ design, and comments, and inspiring conversations regarding the manuscript. We also thank Claire Pelofi, Hao Lu, Maya Pillai, Claire Asenso, Aaron Merlos, and Maiah Xayavong for their assistance in data collection. This research was funded by grants from the National Institutes of Health (R01 DC016119 and U01 AG058532), National Science Foundation (1764010), and the Air Force Office of Scientific Research (FA9550-16-1-0036).

---

**Figure 2–Figure supplement 1.**
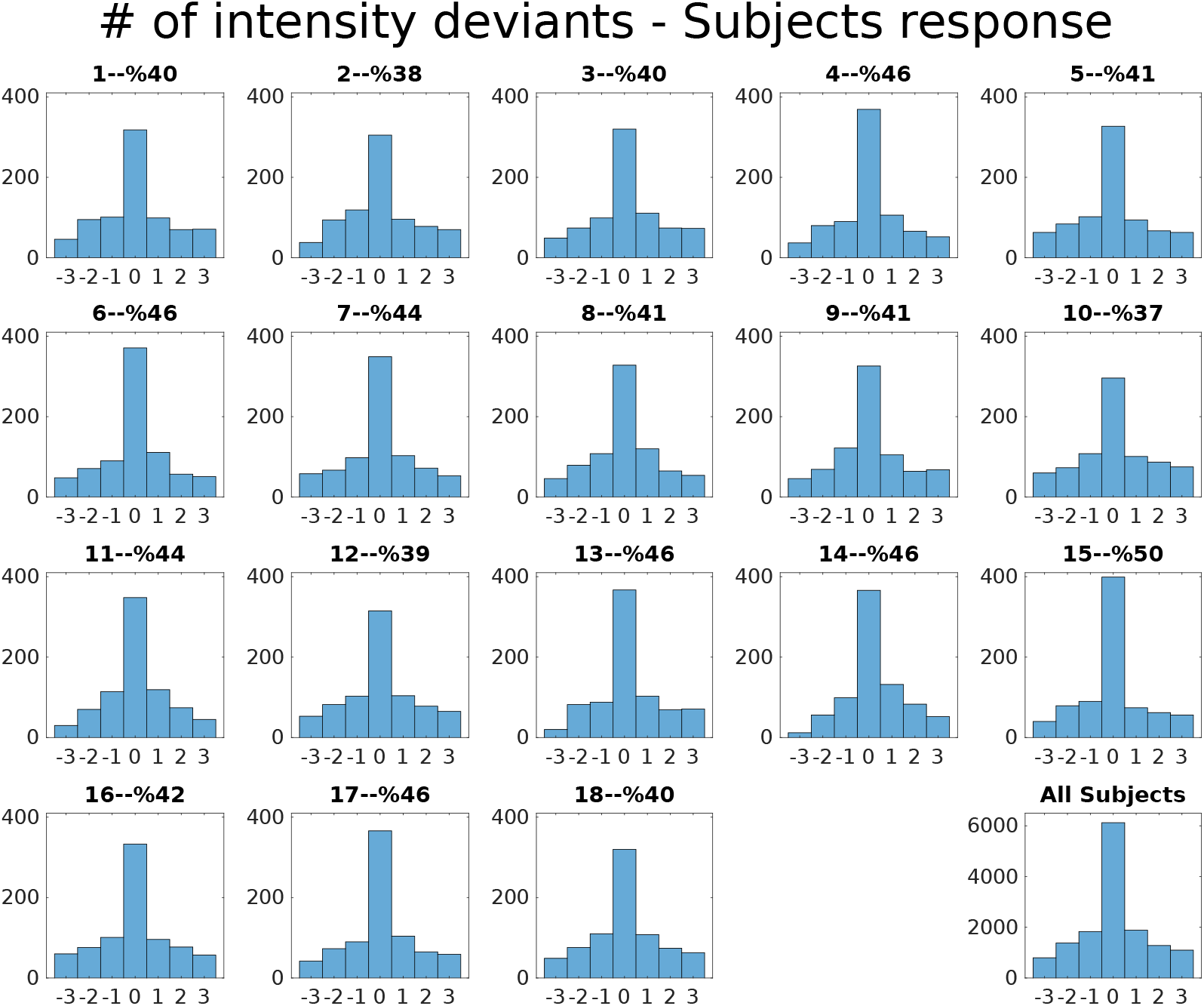
Behavioral Results. In this experiment, listeners were instructed to count the number of deviants in the target (attended) sequence, which was uniformly distributed between 0-3 (four choices) across trials, and hence, the chance level was at %25. Each subplot shows the histogram of the true number of deviants minus the subject’s response. Therefore, in these subplots, “0” means the correct response (hit), positive numbers mean that listeners missed one or some of the deviants, and negatives mean response was larger than the actual number of deviants. Each subplot’s title includes the subject’s number followed by their percentage of correct answers (hit rate). All the subjects performed above the chance level.

**Figure 2–Figure supplement 2.**
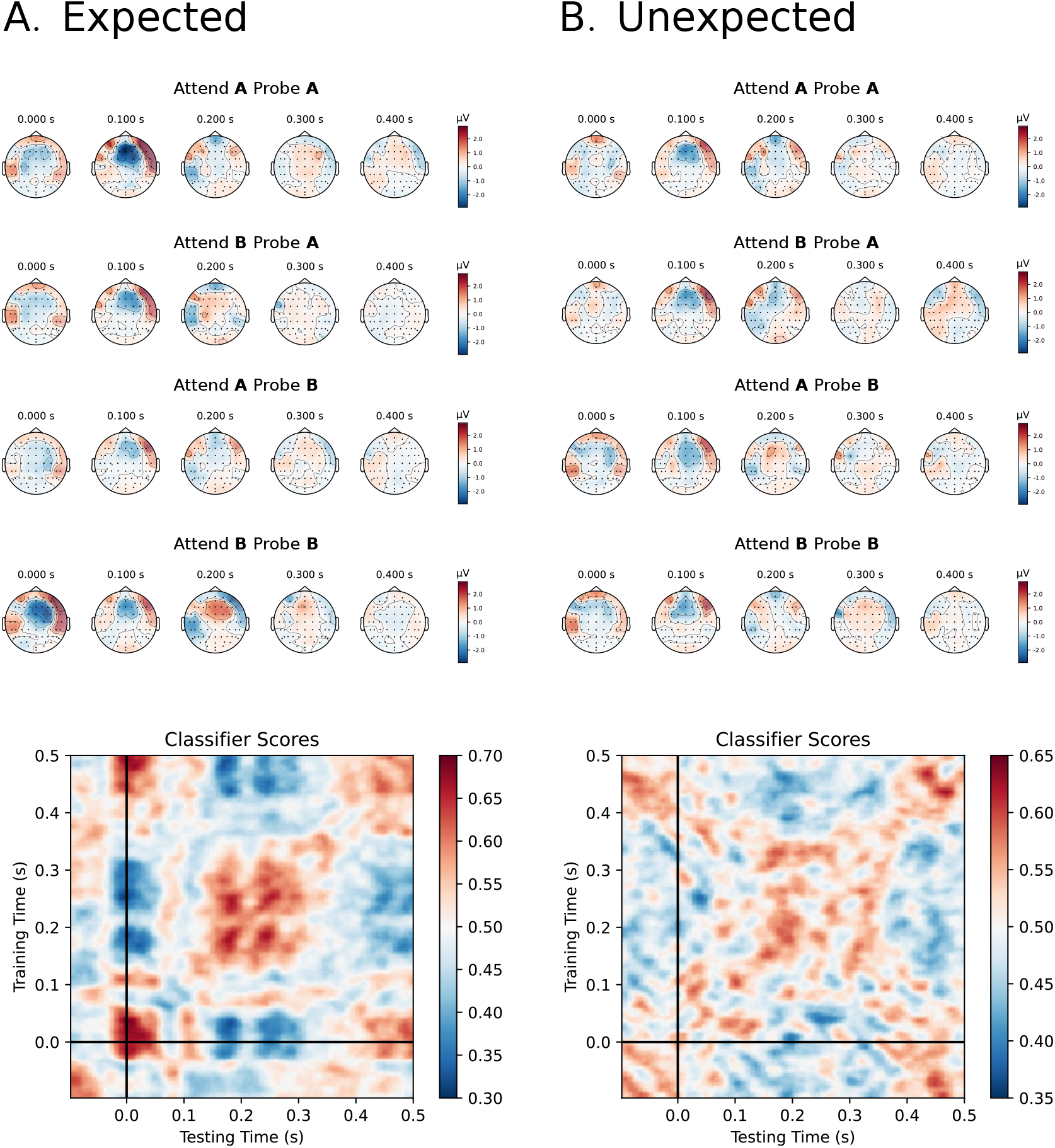
A subject example. Topomaps of probe A and probe B is plotted for different attentional conditions for **(A)** *Expected* and **(B)** *Unexpected* cases. Linear classifiers were trained at each time point on the responses from all 64 channels (topomaps) in order to decode the focus of attention. At the subject level, the trained classifier could capture the differences in the topomap patterns caused by the attentional changes, e.g., the differences between the topomaps of *Attend A probe A* and *Attend B Probe A*. The classifier scores showed the robustness of the effect for a given subject across all trials; in the second-level test (depicted in **Figure 2**B), we showed the robustness of the effect sizes across all subjects.

**Figure 2–Figure supplement 3.**
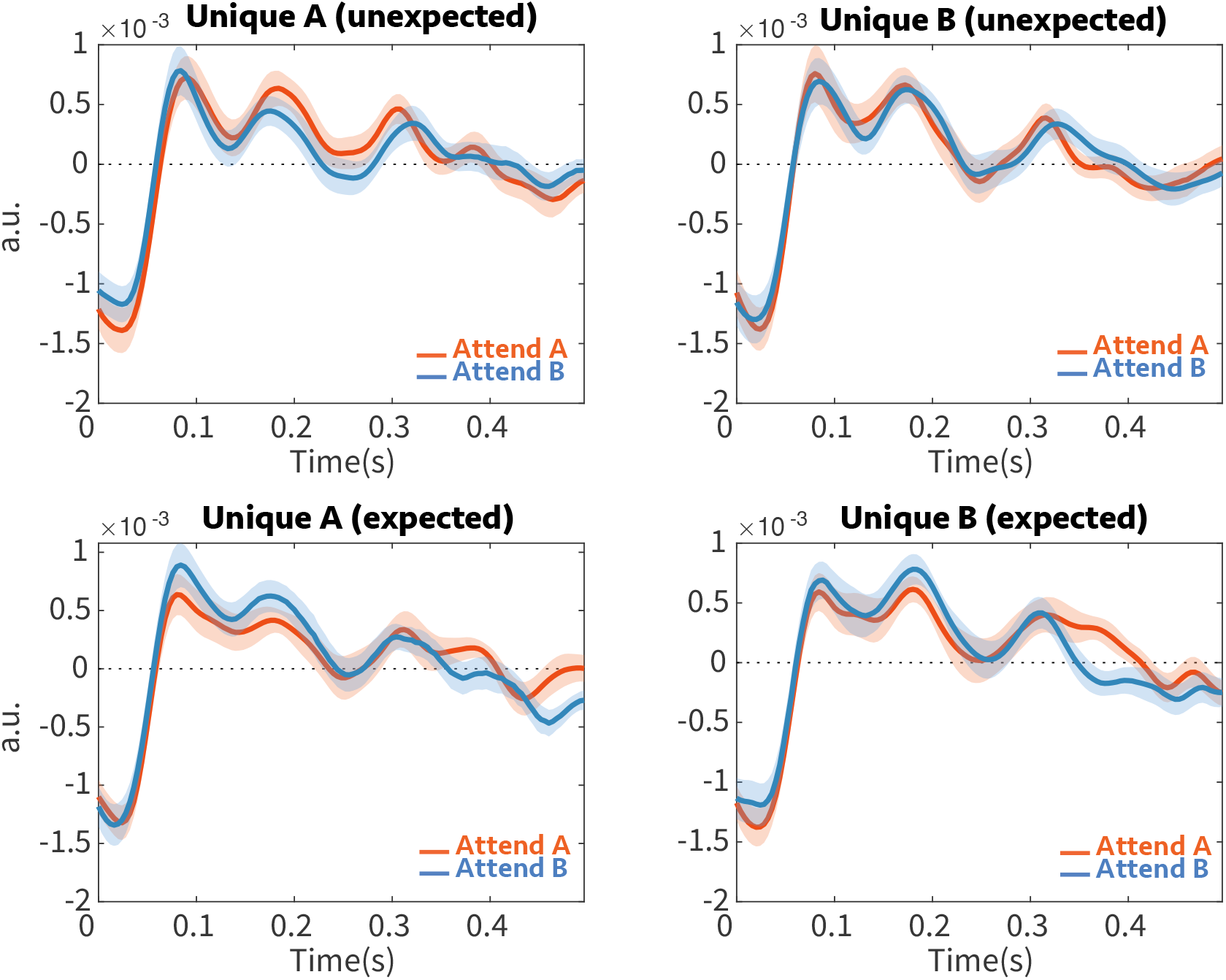
DSS evoked response. The preprocessed EEG data were submitted to the DSS method (see Methods **section**) using the average across trials as a bias filter. The aim was to isolate the most repeatable auditory component by applying a spatial filter. The grand average for the first DSS component is depicted here for probe tones unique to tone complex A (unique A) and tone complex B (unique B) under attend A and attend B conditions, with the onset of the probe at zero. In **Figure 2**C, we subtracted the orange and blue curves in the top-left panel from the same colors on the top-right panel and compared the difference for attend A and attend B. Since attention enhanced the attended probe and suppressed the unattended probe, the probe responses’ difference should had opposite polarity.

**Figure 3–Figure supplement 1.**
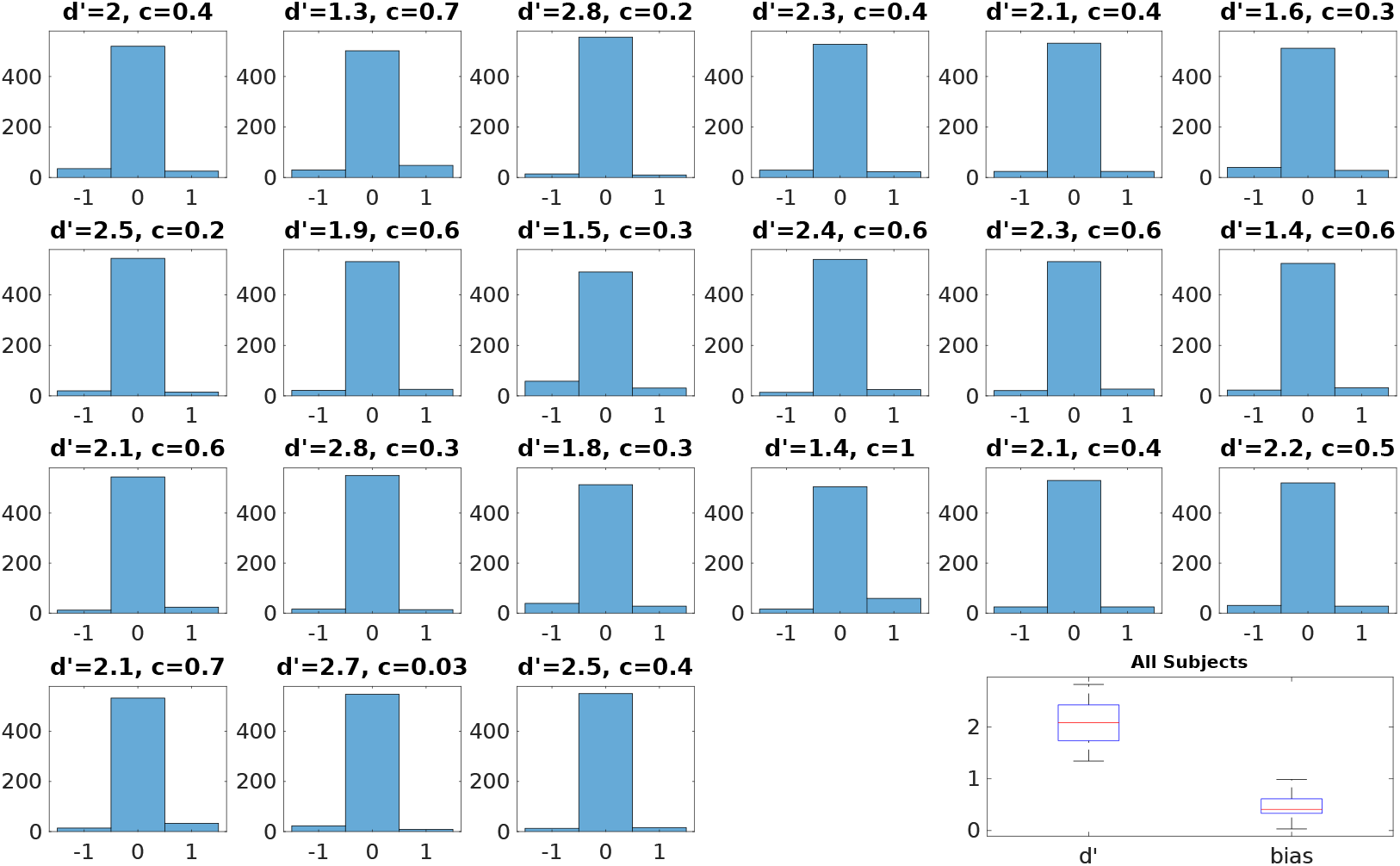
Behavioral Results. In this experiment, listeners were instructed to detect a deviant in the target (attended) sequence. Each subplot shows the histogram of a presence (0 or 1) minus the subject’s response. Therefore, in these subplots, “0” means the correct response (hit), +1 means listeners missed the deviant, and −1 reflects the false alarms. The title of each subplot includes the subject’s d’prime followed by their bias.

**Figure 3–Figure supplement 2.**
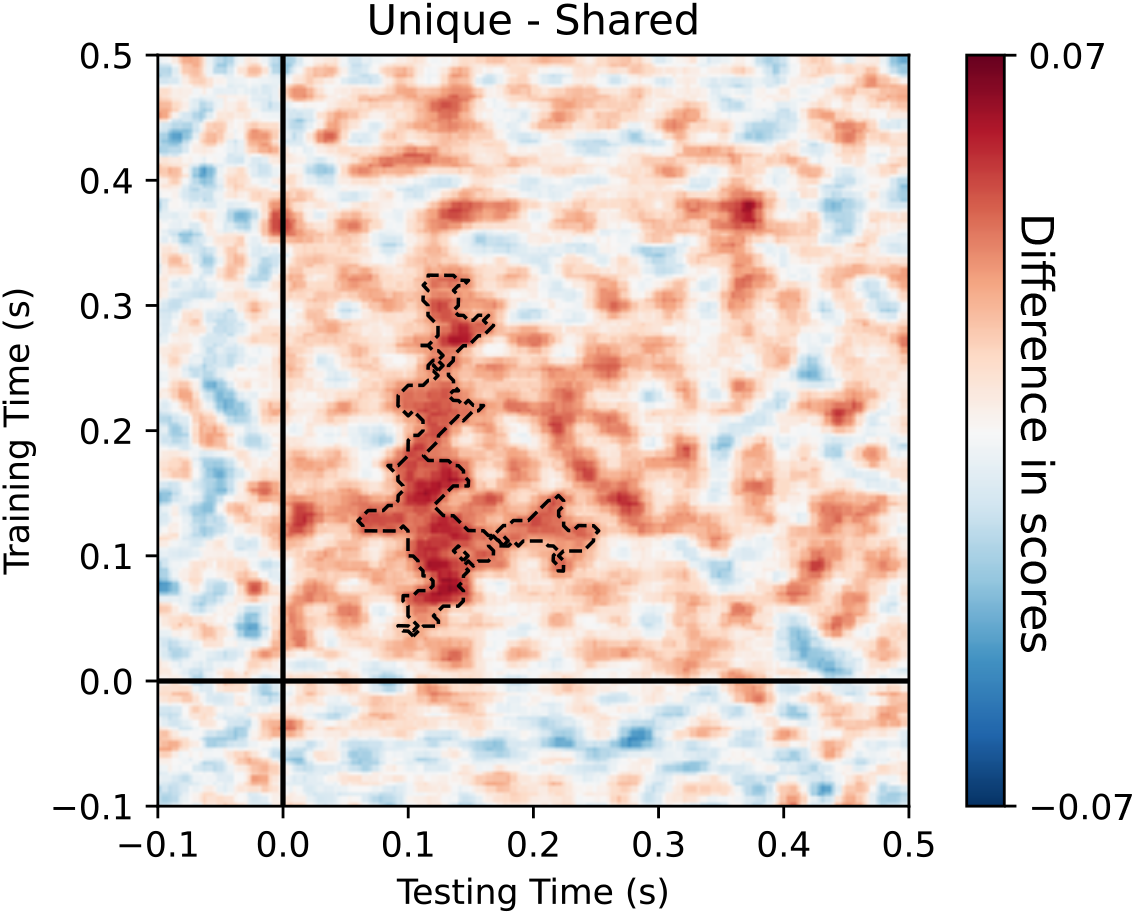
Scores show the average difference between decoding scores of Unique and Shared probe-tone for all subjects. The difference is significant for the time region contoured between the dashed lines (*p* = 0.009).

**Figure 3–Figure supplement 3.**
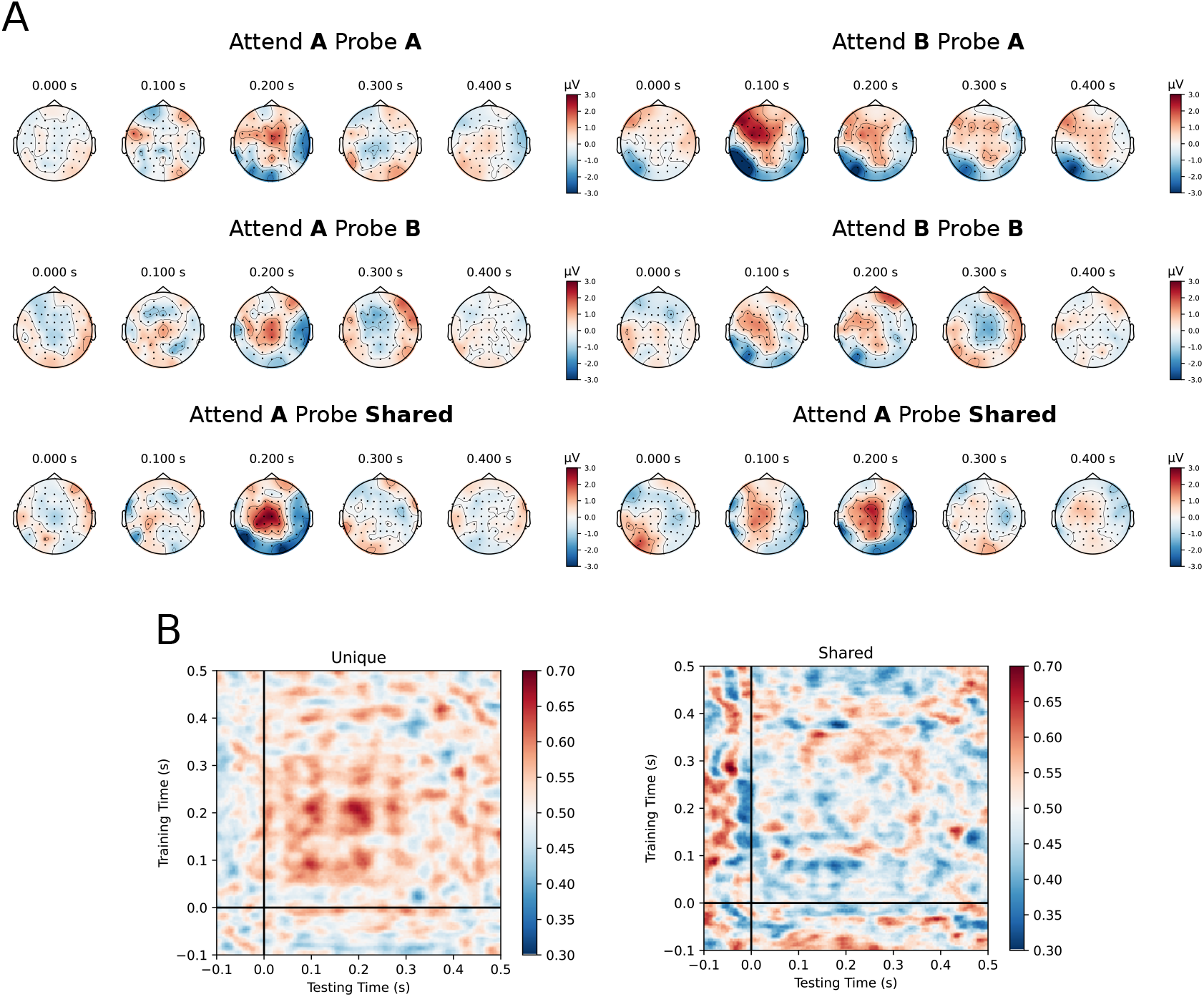
**(A)** Average topomaps of probe A, probe B, and the shared probe were plotted for different attentional conditions. Linear classifiers were trained at each time on the signals from all 64 channels (topomaps) in order to decode the focus of attention. At the subject level, the trained classifier tried to capture the differences in the topomap patterns caused by the attention. **(B)** The classifier scores demonstrated the robustness of the effect for the given subject across all trials. For the unique probe (left), the performance of the classifiers was above chance, which means that there was a consistent difference, i.e., a difference between “*Attend A Probe A”* and “*Attend B Probe A”* in topomap patterns across all trials. Conversely, the shared probe scores suggest that the difference between attentional conditions was not robust since it was not linearly separable.

**Figure 3–Figure supplement 4.**
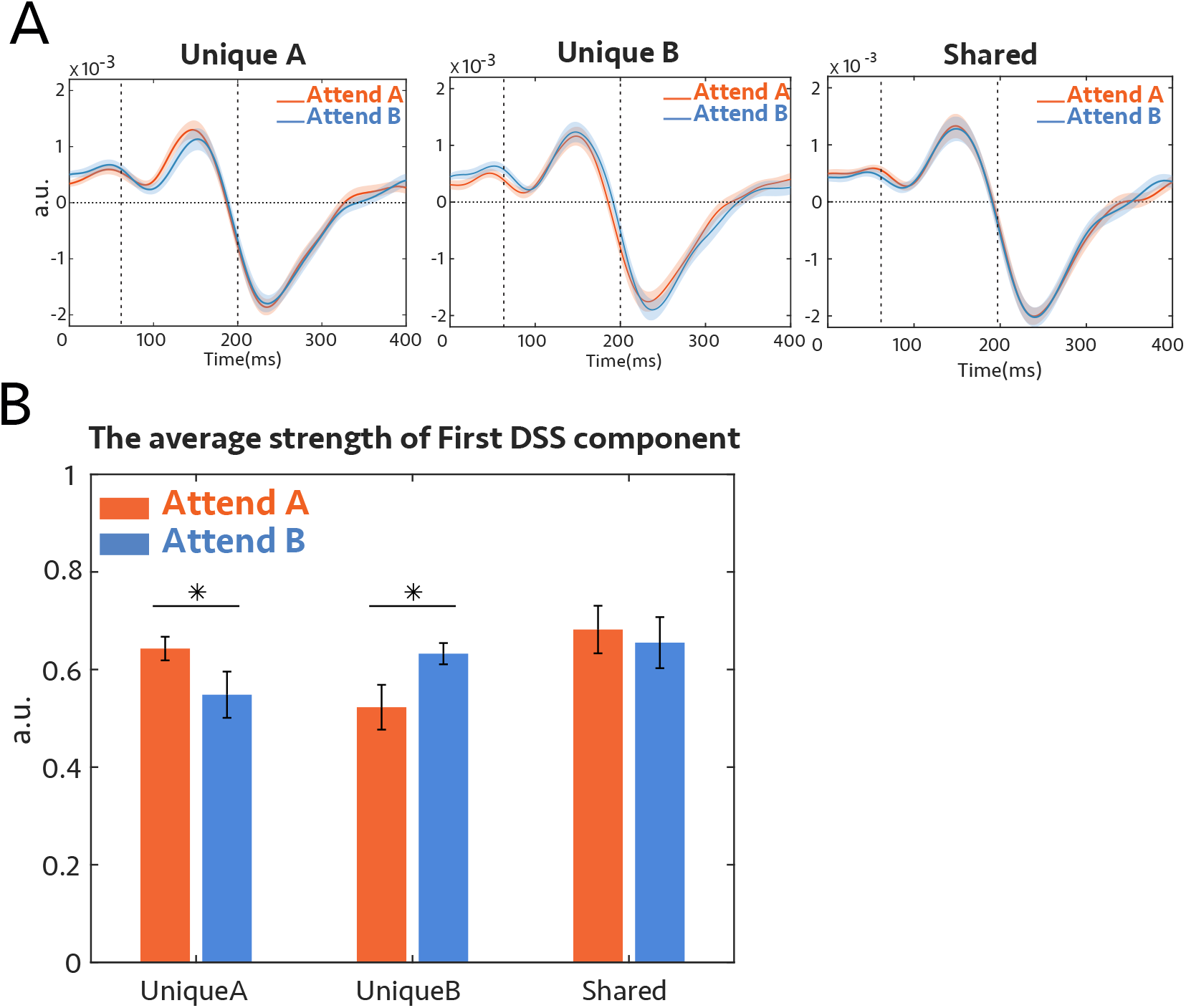
Data were submitted to DSS analysis (see Methods **section**) using the average across trials as a bias filter. The aim was to isolate the most repeatable auditory component by applying a spatial filter. **(A)** Grand average of the most repeatable EEG response to the probe-tone extracted by DSS for each subject; onset of the probe tone is at 0. Left: The response when the probe is at the frequency unique to complex A, middle: when the probe tone is a unique component of complex B, right: The response when the probe tone is a shared component, for attention to tone complex A (orange), and attention to tone complex B (blue). In **Figure 3**C, we subtracted the orange and blue curves in unique A from the same colors in unique B. **(B)** The *average* amplitude of the neural response from 60 ms to 200 ms after the probe-tone onset. For the unique frequency channels, the attended condition has significantly higher power than the unattended condition (*p* = 0.03 for unique A and *p* = 0.01 for unique B), while the *average* of the shared channel does not show any modulation with attention (*p* = 0.6).

**Figure 4–Figure supplement 1.**
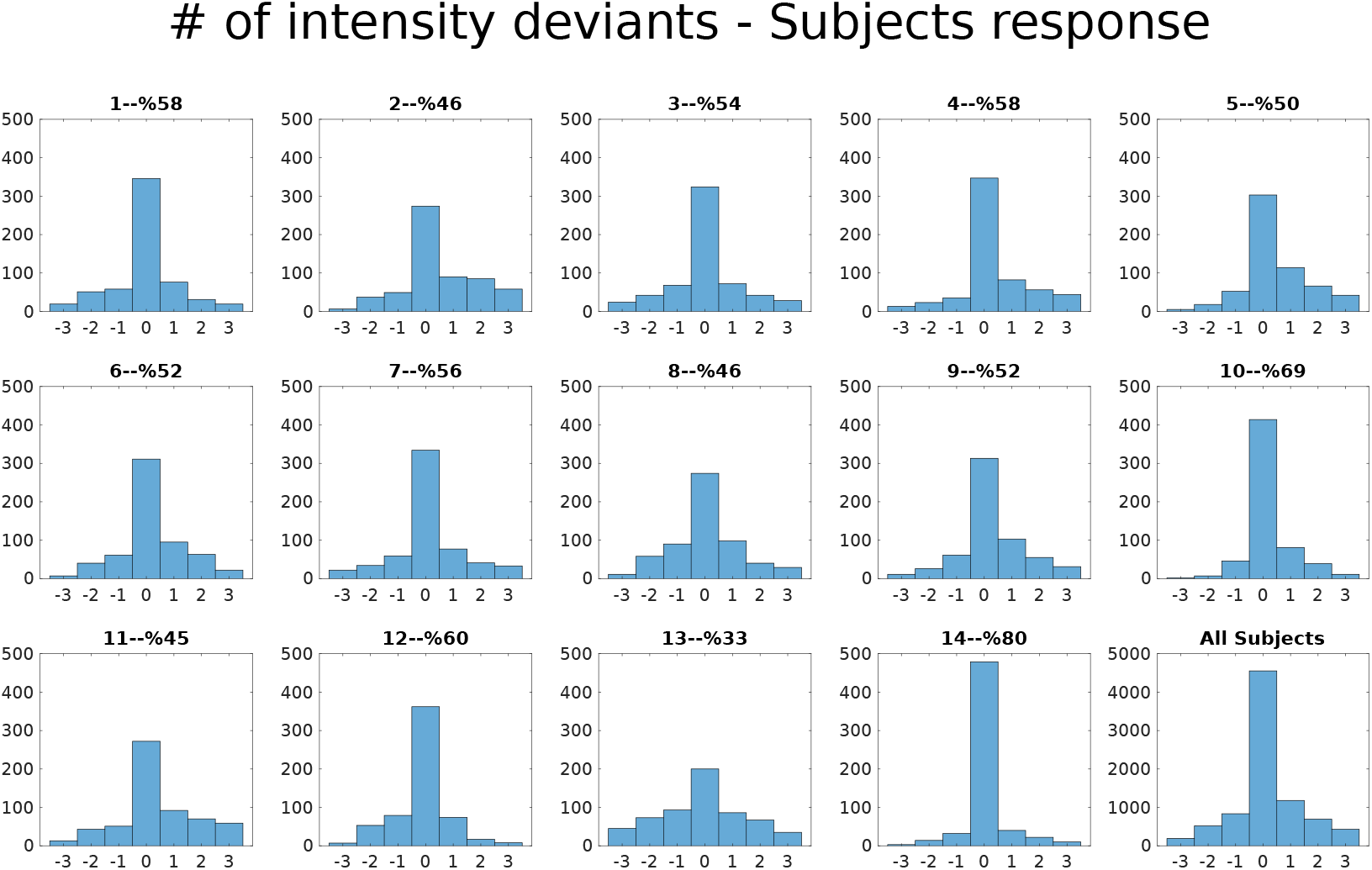
Behavioral Results. In this experiment, listeners were instructed to count the number of deviants in the target (attended) noise sequence, which was uniformly distributed between 0-3 (four choices) across trials, and hence, the chance level was at %25. Each subplot shows the histogram of the true number of deviants minus the subject’s response. Therefore, in these subplots, “0” means the correct response (hit), positive numbers mean that listeners missed one or some of the deviants, and negatives mean response was larger than the actual number of deviants. Each subplot’s title includes the subject’s number followed by their percentage of correct answers (hit rate). All the subjects performed above the chance level (chance level = %25).

**Figure 4–Figure supplement 2.**
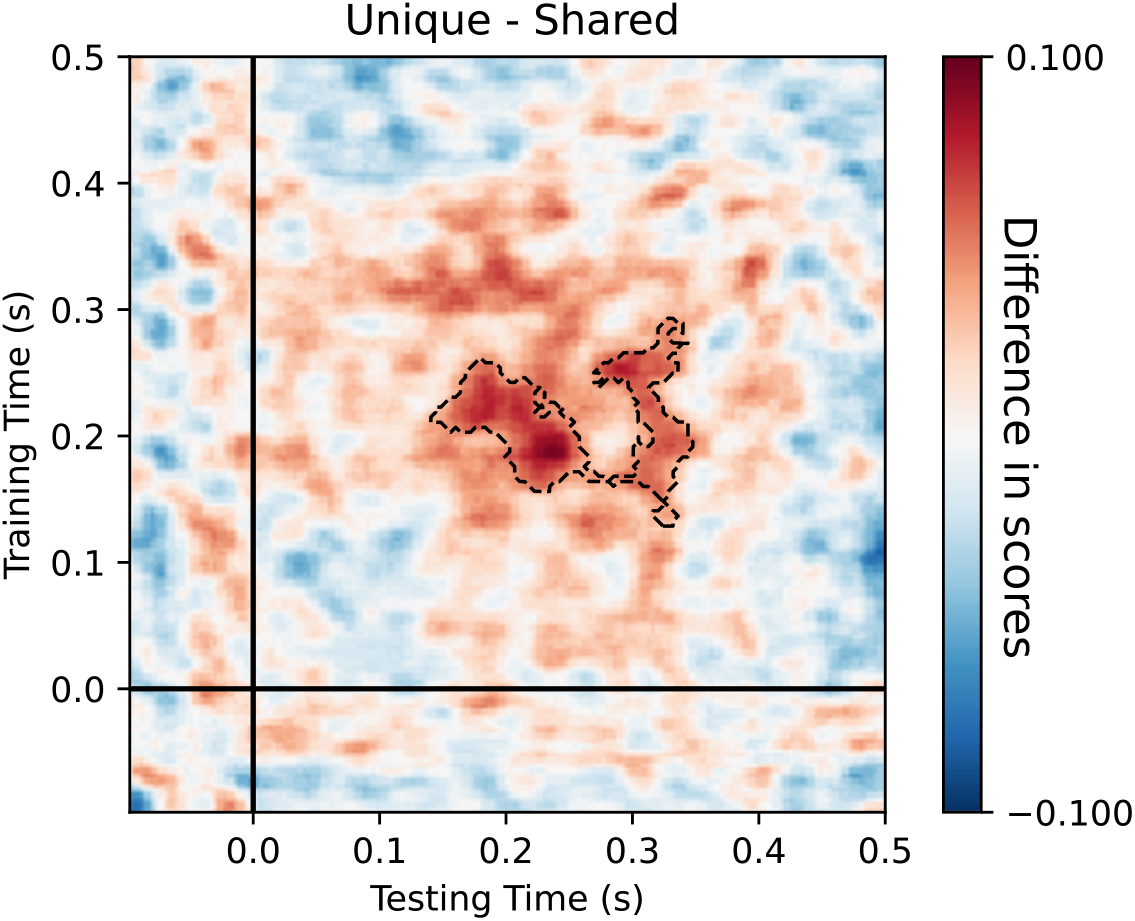
Scores show the average difference between decoding scores of Unique and Shared probe-tone for all subjects. The difference was significant for the time region contoured between the dashed lines (*p* = 0.004).

**Figure 4–Figure supplement 3.**
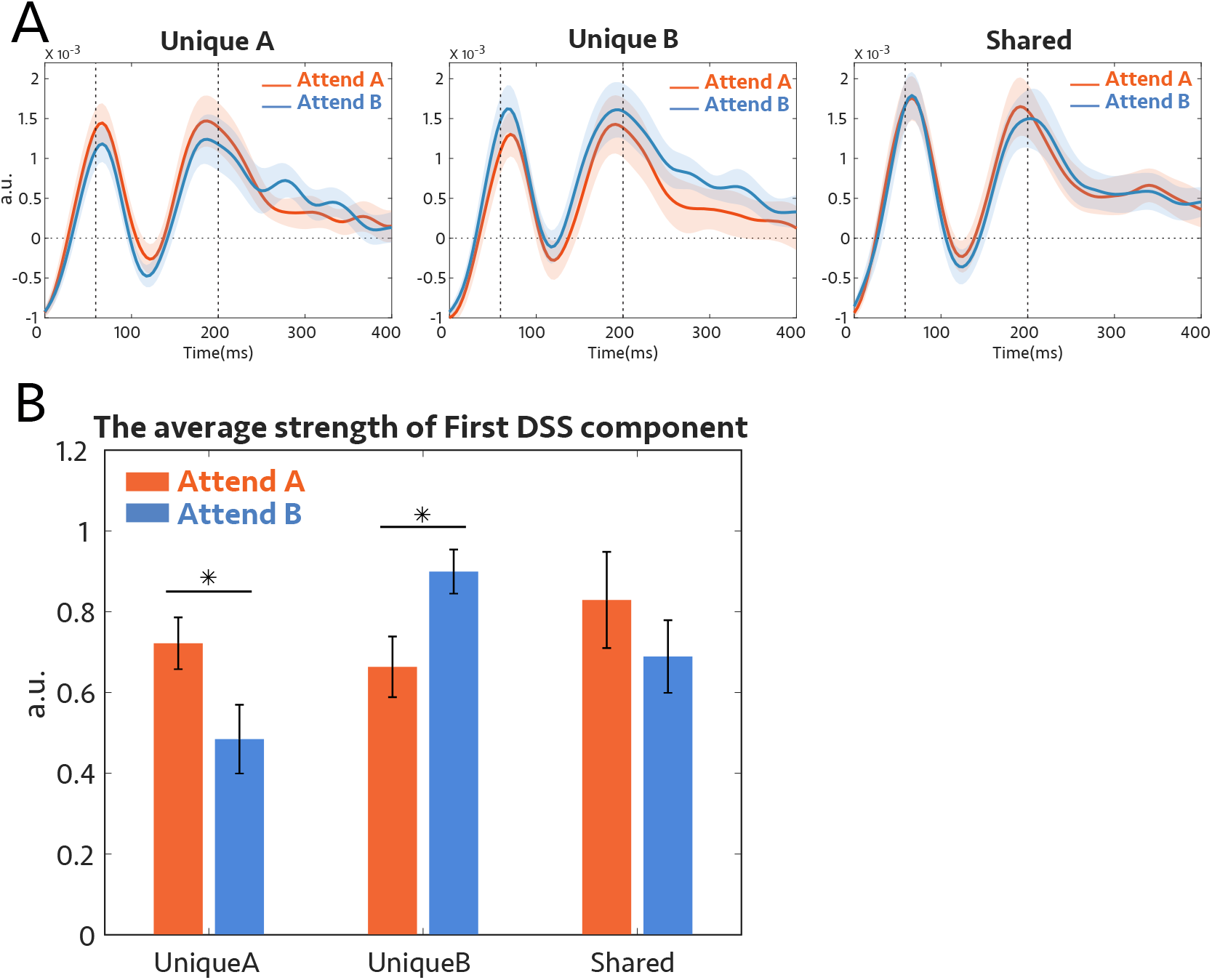
Data were submitted to the DSS using the average across trials as a bias filter. The aim was to isolate the most repeatable auditory component by applying a spatial filter. **(A)** Grand Average of the most repeatable EEG response to the probe-tone extracted by DSS for each subject; onset of the probe tone is at 0. Left: The response when the probe was centered at the unique A frequency channels. Middle: The response to the probe-tone unique to complex B. Right: The response when the probe tone was a shared component, under attend to tone complex A (orange) and attend to tone complex B (blue), the curves are comparable. In **Figure 4**C, we subtracted the orange and blue curves in unique A from the same colors in unique B. **(B)** The average strength of the neural response of the first DSS component from 60 ms to 200 ms after the probe-tone onset. For the unique frequency channels, the attended condition had significantly higher power than the unattended condition (*p* = 0.04 for unique A and *p* = 0.01 for unique B), while the *mean* of the shared channel did not show any modulation with attention (*p* = 0.24).

**Figure 5–Figure supplement 1.**
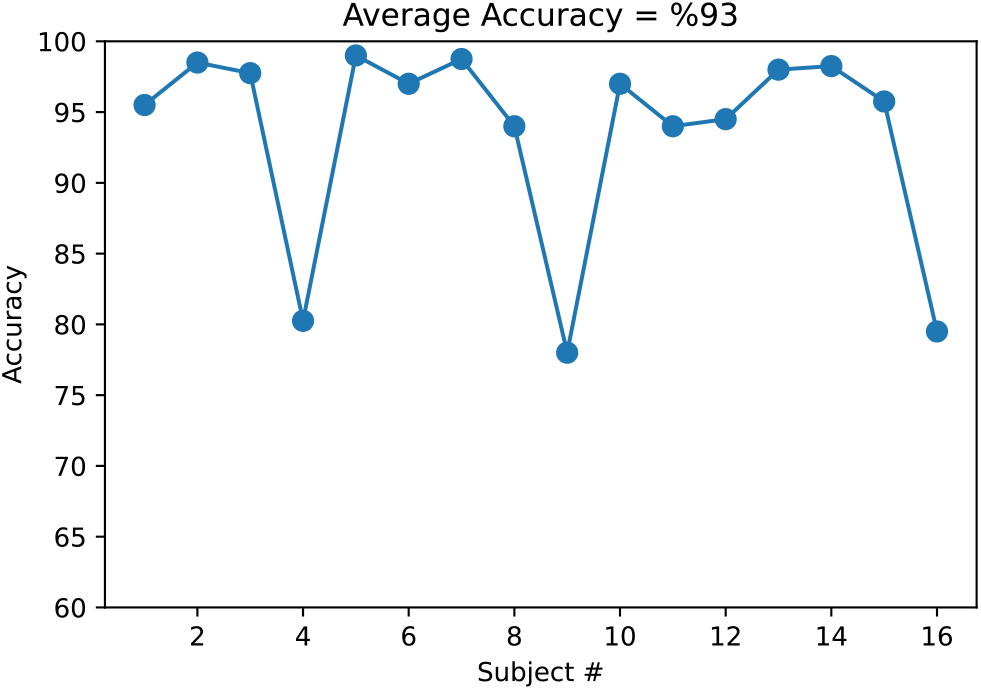
Behavioral Results. In this experiment, listeners were instructed to report the number or the color of the attended speaker. Each point shows the accuracy for each subject.

**Figure 5–Figure supplement 2.**
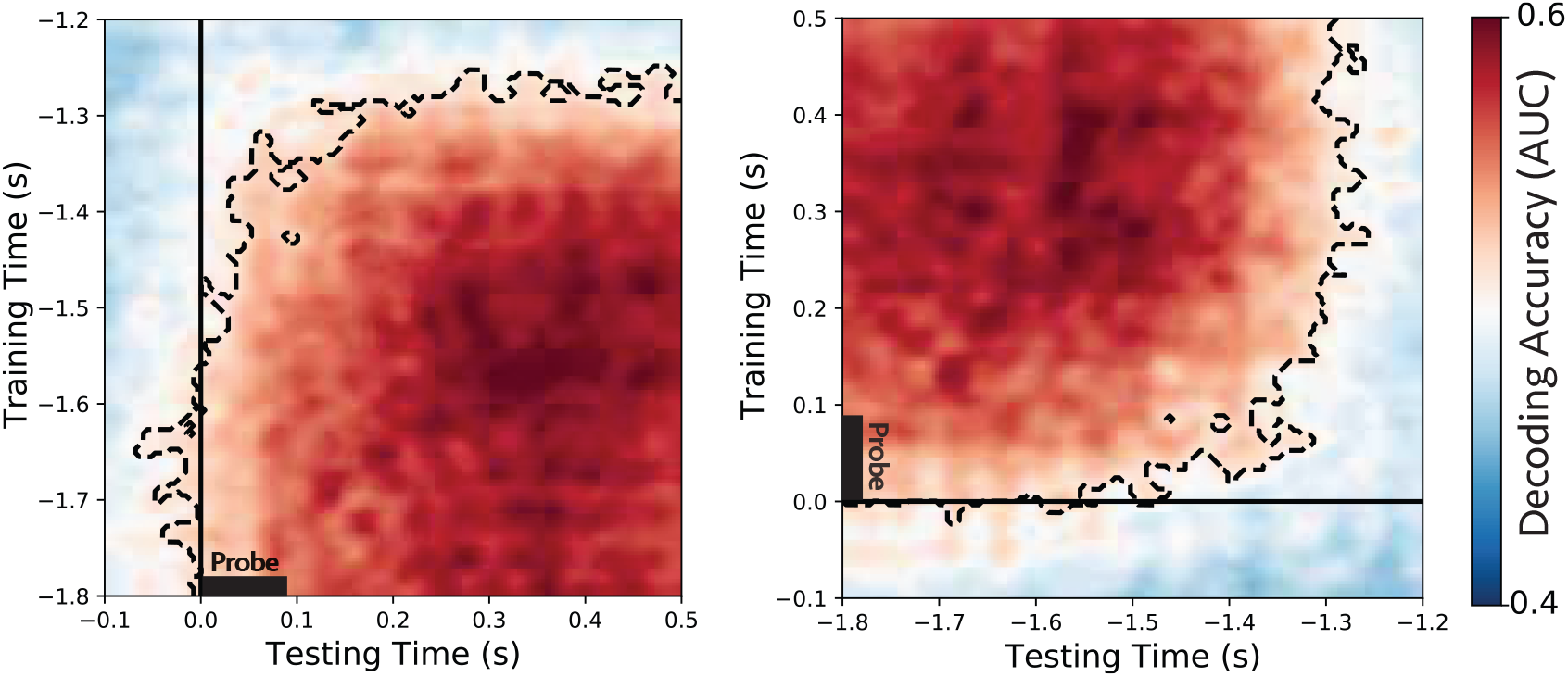
Generalizing decoders across time. Classifiers trained and tested separately at each time instant in two other 600 ms time windows. *Left:* trained at the beginning of the speech (−1.8 sec to −1.2 sec) and tested during the probe-tone (−100ms to 500ms). *Middle:* trained during the probe time window (−0.1 sec to 0.5 sec) and tested during the beginning of the speech (−1.8 sec to −1.2 sec). These results suggest that the modulatory effect of attention is generalizable across times during speech and probe-tone.

**Figure 5–Figure supplement 3.** Evoked responses at channel Cz

**Figure 6–Figure supplement 1.** Behavioral results.

**Figure 6–Figure supplement 2.** Evoked responses

## References

Atilgan H, Bizley JK. Training enhances the ability of listeners to exploit visual information for auditory scene analysis. Cognition. 2021 mar; 208. https://pubmed.ncbi.nlm.nih.gov/33373937/, doi: 10.1016/j.cognition.2020.104529.

Atilgan H, Town S, Wood K, Jones G, Maddox R, Lee AK, Bizley J. Integration of Visual Information in Auditory Cortex Promotes Auditory Scene Analysis through Multisensory Binding. Neuron. 2018 01; 97:640–655. doi: 10.1016/j.neuron.2017.12.034.

Bernstein L, Auer E, Takayanagi S. Auditory speech detection in noise enhanced by lipreading. Speech Communication. 2004 10; 44:5–18. doi: 10.1016/j.specom.2004.10.011.

Bizley JK, Maddox RK, Lee AKC. Defining Auditory-Visual Objects: Behavioral Tests and Physiological Mechanisms. Trends in Neurosciences. 2016 feb; 39(2):74–85. /pmc/articles/PMC4738154//pmc/articles/PMC4738154/?report=abstract https://www.ncbi.nlm.nih.gov/pmc/articles/PMC4738154/, doi: 10.1016/j.tins.2015.12.007.

Bolia R, Nelson W, Ericson M, Simpson B. A speech corpus for multitalker communications research. The Journal of the Acoustical Society of America. 2000 03; 107:1065–6. doi: 10.1121/1.428288.

Brainard DH. The Psychophysics Toolbox. Spatial Vision. 1997; 10(4):433–436. https://brill.com/view/journals/sv/10/4/article-p433_15.xml, doi: https://doi.org/10.1163/156856897X00357.

Bregman AS, Liao C, Levitan R. Auditory grouping based on fundamental frequency and formant peak frequency. Canadian journal of psychology. 1990; 44(3):400–413. /record/1991-06393-001, doi: 10.1037/h0084255.

Bregman A. Auditory Scene Analysis: The Perceptual Organization of Sound. Journal of The Acoustical Society of America - J ACOUST SOC AMER. 1990 01; 95:250. doi: 10.1121/1.408434.

Bregman AS, Campbell J. Primary auditory stream segregation and perception of order in rapid sequences of tones. Journal of Experimental Psychology. 1971; 89(2):244–249. http://doi.apa.org/getdoi.cfm?doi=10.1037/h0031163, doi: 10.1037/h0031163.

Brungart DS, Simpson BD, Ericson MA, Scott KR. Informational and energetic masking effects in the perception of multiple simultaneous talkers. The Journal of the Acoustical Society of America. 2001; 110(5):2527–2538. https://doi.org/10.1121/1.1408946, doi: 10.1121/1.1408946.

Carlyon R, Thompson S, Heinrich A, Pulvermüller F, Davis M, Shtyrov Y, Cusack R, Johnsrude I. 47. In: Objective Measures of Auditory Scene Analysis Springer, New York, NY; 2010. p. 507–519. doi: 10.1007/978-1-4419-5686-6_47.

de Cheveigné A, Arzounian D. Robust detrending, rereferencing, outlier detection, and inpainting for multichannel data. NeuroImage. 2018 may; 172:903–912. https://www.sciencedirect.com/science/article/pii/S1053811918300351, doi: 10.1016/J.NEUROIMAGE.2018.01.035.

de Cheveigné A, Simon JZ. Denoising based on time-shift PCA. Journal of Neuroscience Methods. 2007 sep; 165(2):297–305. doi: 10.1016/j.jneumeth.2007.06.003.

Choi I, Rajaram S, Varghese L, Shinn-Cunningham B. Quantifying attentional modulation of auditory-evoked cortical responses from single-trial electroencephalography. Frontiers in human neuroscience. 2013 04; 7:115. doi: 10.3389/fnhum.2013.00115.

Crosse MJ, Di Liberto GM, Lalor EC. Eye Can Hear Clearly Now: Inverse Effectiveness in Natural Audiovisual Speech Processing Relies on Long-Term Crossmodal Temporal Integration. Journal of Neuroscience. 2016; 36(38):9888–9895. https://www.jneurosci.org/content/36/38/9888, doi: 10.1523/JNEUROSCI.1396-16.2016.

De Cheveigné A, Parra LC. Joint decorrelation, a versatile tool for multichannel data analysis. NeuroImage. 2014; 98:487–505. http://dx.doi.org/10.1016/j.neuroimage.2014.05.068, doi: 10.1016/j.neuroimage.2014.05.068.

De Cheveigné A, Simon JZ. Denoising based on spatial filtering. Journal of Neuroscience Methods. 2008; 171(2):331–339. doi: 10.1016/j.jneumeth.2008.03.015.

Ding N, Simon JZ. Emergence of neural encoding of auditory objects while listening to competing speakers. Proceedings of the National Academy of Sciences. 2012; 109(29):11854–11859. https://www.pnas.org/content/109/29/11854, doi: 10.1073/pnas.1205381109.

Elhilali M, Ma L, Micheyl C, Oxenham AJ, Shamma SA. Temporal Coherence in the Perceptual Organization and Cortical Representation of Auditory Scenes. Neuron. 2009; 61(2):317–329. http://dx.doi.org/10.1016/j.neuron.2008.12.005, doi: 10.1016/j.neuron.2008.12.005.

Gramfort A, Luessi M, Larson E, Engemann D, Strohmeier D, Brodbeck C, Goj R, Jas M, Brooks T, Parkkonen L, Hämäläinen M. MEG and EEG data analysis with MNE-Python. Frontiers in Neuroscience. 2013; 7:267. https://www.frontiersin.org/article/10.3389/fnins.2013.00267, doi: 10.3389/fnins.2013.00267.

Grimault N, Micheyl C, Carlyon RP, Arthaud P, Collet L. Perceptual auditory stream segregation of sequences of complex sounds in subjects with normal and impaired hearing. British journal of audiology. 2001 07; 35:173–82. doi: 10.1080/00305364.2001.11745235.

King JR, Dehaene S. Characterizing the dynamics of mental representations: the temporal generalization method. Trends in Cognitive Sciences. 2014 apr; 18(4):203–210. https://www.sciencedirect.com/science/article/pii/S1364661314000199, doi: 10.1016/J.TICS.2014.01.002.

King JR, Pescetelli N, Dehaene S. Brain Mechanisms Underlying the Brief Maintenance of Seen and Unseen Sensory Information. Neuron. 2016 12; 92:1122–1134. doi: 10.1016/j.neuron.2016.10.051.

Kleiner M, Brainard DH, Pelli DG, Broussard C, Wolf T, Niehorster D. What’s new in Psychtoolbox-3? A free cross-platform toolkit for psychophysiscs with Matlab and GNU/Octave; 2007.

Krishnan L, Elhilali M, Shamma S. Segregating Complex Sound Sources through Temporal Coherence. PLoS Computational Biology. 2014; 10(12). doi: 10.1371/journal.pcbi.1003985.

Krogholt Christiansen S, Oxenham AJ. Assessing the effects of temporal coherence on auditory stream formation through comodulation masking release. The Journal of the Acoustical Society of America. 2014; 135(6):3520–3529. https://www.ncbi.nlm.nih.gov/pmc/articles/PMC4048442/, doi: 10.1121/1.4872300.

Lee SH, Blake R. Visual Form Created Soley from Temporal Structure. Science (New York, NY). 1999 06; 284:1165–8. doi: 10.1126/science.284.5417.1165.

Lu K, Xu Y, Yin P, Oxenham AJ, Fritz JB, Shamma SA. Temporal coherence structure rapidly shapes neuronal interactions. Nature Communications. 2017; 8:13900. http://www.nature.com/doifinder/10.1038/ncomms13900, doi: 10.1038/ncomms13900.

Maris E, Oostenveld R. Nonparametric statistical testing of EEG- and MEG-data. Journal of Neuroscience Methods. 2007; 164(1):177–190. http://www.sciencedirect.com/science/article/pii/S0165027007001707, doi: https://doi.org/10.1016/j.jneumeth.2007.03.024.

Mesgarani N, Chang EF. Selective cortical representation of attended speaker in multi-talker speech perception. Nature. 2012; 485(7397):233–6. http://www.pubmedcentral.nih.gov/articlerender.fcgi?artid=3870007{&}tool=pmcentrez{&}rendertype=abstract, doi: 10.1038/nature11020.

Micheyl C, Hanson C, Demany L, Oxenham AJ, Shamma S. Auditory stream segregation for alternating and synchronous tones. Journal of Experimental Psychology: Human Perception and Performance. 2013; 39(6):1568–1580. https://pubmed.ncbi.nlm.nih.gov/23544676/, doi: 10.1037/a0032241.

Micheyl C, Kreft H, Shamma S, Oxenham AJ. Temporal coherence versus harmonicity in auditory stream formation. The Journal of the Acoustical Society of America. 2013; 133(3):EL188–EL194. https://www.ncbi.nlm.nih.gov/pmc/articles/PMC3579859/, doi: 10.1121/1.4789866.

Micheyl C, Oxenham AJ. Objective and subjective psychophysical measures of auditory stream integration and segregation. JARO - Journal of the Association for Research in Otolaryngology. 2010; 11(4):709–724. http://www.tc.umn.edu/{~}cmicheyl/demos.html., doi: 10.1007/s10162-010-0227-2.

Middlebrooks JC, Simon JZ, Popper AN, Editors RRF. The Auditory System at the Cocktail Party, vol. 60. Springer Handbook of Auditory research; 2017. http://link.springer.com/10.1007/978-3-319-51662-2, doi: 10.1007/978-3-319-51662-2.

Moore B, Gockel H. Factors Influencing Sequential Stream Segregation. Acta Acustica united with Acustica. 2002 05; 88:320–333.

O’Sullivan AE, Crosse MJ, Di Liberto GM, de Cheveigné A, Lalor EC. Neurophysiological indices of audiovisual speech integration are enhanced at the phonetic level for speech in noise. bioRxiv. 2020; https://www.biorxiv.org/content/early/2020/04/20/2020.04.18.048124, doi: 10.1101/2020.04.18.048124.

O’Sullivan J, Herrero J, Smith E, Schevon C, McKhann GM, Sheth SA, Mehta AD, Mesgarani N. Hierarchical Encoding of Attended Auditory Objects in Multi-talker Speech Perception. Neuron. 2019; 104(6):1195–1209.e3. https://doi.org/10.1016/j.neuron.2019.09.007, doi: 10.1016/j.neuron.2019.09.007.

O’Sullivan J, Shamma SA, Lalor EC. Evidence for Neural Computations of Temporal Coherence in an Auditory Scene and Their Enhancement during Active Listening. Journal of Neuroscience. 2015; 35(18):7256–7263. https://www.jneurosci.org/content/35/18/7256, doi: 10.1523/JNEUROSCI.4973-14.2015.

Pedregosa F, Varoquaux G, Gramfort A, Michel V, Thirion B, Grisel O, Blondel M, Prettenhofer P, Weiss R, Dubourg V, Vanderplas J, Passos A, Cournapeau D, Brucher M, Perrot M, Édouard Duchesnay. Scikit- learn: Machine Learning in Python. Journal of Machine Learning Research. 2011; 12(85):2825–2830. http://jmlr.org/papers/v12/pedregosa11a.html.

Pelli DG. The VideoToolbox software for visual psychophysics: transforming numbers into movies. Spatial Vision. 1997; 10(4):437–442. https://brill.com/view/journals/sv/10/4/article-p437_16.xml, doi: https://doi.org/10.1163/156856897X00366.

Pinheiro-Chagas P, Piazza M, Dehaene S. Decoding the processing stages of mental arithmetic with magnetoencephalography. Cortex. 2018 jul; https://www.sciencedirect.com/science/article/pii/S0010945218302351, doi: 10.1016/J.CORTEX.2018.07.018.

Power A, Foxe J, Forde EJ, Reilly R, Lalor E. At what time is the cocktail party? A late locus of selective attention to natural speech. The European journal of neuroscience. 2012 03; 35:1497–503. doi: 10.1111/j.1460-9568.2012.08060.x.

Shamma S, Elhilali M. 2. In: Fritzsch B, editor. Chapter: Binding, Scene Analysis and Higher Cortical Centers in The Senses: A Comprehensive Reference, 2nd Edition ELSEVIER ACADEMIC Press; 2020. https://books.google.com/books?id=kLNWzQEACAAJ.

Shamma SA, Elhilali M, Micheyl C. Temporal coherence and attention in auditory scene analysis. Trends in Neurosciences. 2011; 34(3):114–123. http://dx.doi.org/10.1016/j.tins.2010.11.002, doi: 10.1016/j.tins.2010.11.002.

Singh PG. Perceptual organization of complex-tone sequences: A tradeoff between pitch and timbre. Journal of the Acoustical Society of America. 1987; 82(3):886–899. https://pubmed.ncbi.nlm.nih.gov/3655122/, doi: 10.1121/1.395287.

Snyder J, Alain C, Picton T. Effects of Attention on Neuroelectric Correlates of Auditory Stream Segregation. Journal of cognitive neuroscience. 2006 02; 18:1–13. doi: 10.1162/089892906775250021.

Stokes M, Wolff M, Spaak E. Decoding Rich Spatial Information with High Temporal Resolution. Trends in Cognitive Sciences. 2015 11; 19:636–638. doi: 10.1016/j.tics.2015.08.016.

Sussman ES. Auditory Scene Analysis: An Attention Perspective. Journal of Speech, Language, and Hearing Research. 2017; 60(10):2989–3000. https://pubs.asha.org/doi/abs/10.1044/2017_JSLHR-H-17-0041, doi: 10.1044/2017\_JSLHR-H-17-0041.

Teki S, Barascud N, Picard S, Payne C, Griffiths TD, Chait M. Neural Correlates of Auditory Figure-Ground Segregation Based on Temporal Coherence. Cerebral Cortex (New York, NY). 2016; 26(9):3669. https://www.ncbi.nlm.nih.gov/pmc/articles/PMC5004755/, doi: 10.1093/CERCOR/BHW173.

Teki S, Chait M, Kumar S, Shamma S, Griffiths TD. Segregation of complex acoustic scenes based on temporal coherence. eLife. 2013 jul; 2013(2). https://elifesciences.org/articles/00699, doi: 10.7554/eLife.00699.

Wilcoxon F. Individual Comparisons by Ranking Methods. Biometrics Bulletin. 1945; 1(6):80–83. http://www.jstor.org/stable/3001968.

Wolff M, Jochim J, Akyürek E, Stokes M. Dynamic hidden states underlying working-memory-guided behavior. Nature Neuroscience. 2017 04; 20. doi: 10.1038/nn.4546.

Xiang J, Simon J, Elhilali M. Competing Streams at the Cocktail Party: Exploring the Mechanisms of Attention and Temporal Integration. Journal of Neuroscience. 2010; 30(36):12084–12093. https://www.jneurosci.org/content/30/36/12084, doi: 10.1523/JNEUROSCI.0827-10.2010.

Zera J, Green DM. Effect of signal component phase on asynchrony discrimination. The Journal of the Acoustical Society of America. 1995; 98(2):817–827. https://doi.org/10.1121/1.413508, doi: 10.1121/1.413508.

Zera S. Detectisoqahng temporal onset and offset asynchrony in multicomponent complexes. Journal of the Acoustical Society of America. 1993; 93(2):1038–1052. https://pubmed.ncbi.nlm.nih.gov/8445115/, doi: 10.1121/1.405552.

